# Genome-wide profiling of RNA 2’-*O*-methylation in neurons and identification of orphan snoRNA targets

**DOI:** 10.64898/2025.12.17.694928

**Authors:** Xuan Ye, Yaling Liu, Yinzhou Zhu, Sarah A. Alshawi, Brittany A. Elliott, Christopher L. Holley, Jean-Denis Beaudoin, Brenton R. Graveley, Gordon G. Carmichael

## Abstract

We have compared genome-wide patterns of RNA 2’-*O*-methylation (Nm) between two isogenic pairs of neurons. Each pair includes one line harboring a small deletion of orphan box C/D snoRNAs (SNORD116s) from the paternal chr15q11-q13 region. One isogenic pair also differs in expression of SNORD113/114 snoRNAs from chr14q32.2. Wild-type and modified cells were differentiated into cortical neurons, and genome-wide patterns of Nm identified. Neurons display a distinctive signature of rRNA modification compared to undifferentiated stem cells. We further identified thousands of shared Nm sites in mRNAs, lncRNAs and small RNAs. Most sites do not exhibit canonical complementarity to snoRNAs, but a number exhibit strong complementarity to U3 snoRNA, not previously shown to direct Nm. Evidence from cross-linking and sequencing of hybrids (CLASH) suggests that U3 is proximally associated with a subset of 2’-*O*-methylation events. Finally, we identify a number of apparent canonical targets of SNORD113, SNORD114 and SNORD116 snoRNAs. These data present a comprehensive characterization of the Nm landscape in neurons and, for the first time, allow the assignment of Nm sites targeted by specific orphan snoRNAs associated with neurodevelopmental and other disorders.

## INTRODUCTION

To date, over 170 distinct RNA modifications have been identified and characterized^1–3^. These modifications, which don’t alter the RNA sequence, play important roles in regulating gene expression and various physiological and pathological processes. Among these RNA modifications, 2’-*O*-methylation (Nm) is a widespread RNA modification found in multiple RNA types and species,^4^ and it is among the most abundant in the cell owing to numerous sites on rRNAs and small RNAs. 2’-*O*-methylation in RNA is thought to primarily function to increase RNA stability, promote RNA folding, facilitate protein binding, and make RNA less susceptible to hydrolysis and cleavage^5^.

2’-*O*-methylation is especially important in ribosome assembly, maturation, and function in translation^6^. Emerging research indicates that diverse rRNA methylation patterns can precisely modulate ribosome activity, enabling specialized translation tailored to various cellular environments. This adaptability of rRNA methylation underscores its importance in regulating gene expression, demonstrating that it’s not merely a structural element but a dynamic process impacting cellular protein production^7^. Thus, some rRNA sites are modified to different levels in different cells^8–12^. Mechanistically, Nm stabilizes RNA structures^13^ and confers resistance to nucleophilic attack and 3’–5’ degradation^8^. Other transcripts, including small RNAs, viral RNAs, and mRNAs, have been reported to also harbor Nm residues^14–22^. Nm can regulate RNA metabolism by influencing processing events such as splicing or stability and by modulating translation efficiency and fidelity^23–25^. In mRNA, Nm has been reported to inhibit translation by disrupting tRNA decoding^16,17^ and by enhancing RNA stability^20^. Furthermore, Nm modifications can disrupt long-range RNA tertiary interactions that depend on the 2’-hydroxyl for hydrogen bonding and metal ion coordination^26^ and the presence of an Nm can sterically hinder RNA-binding protein association^27,28^. Through these multifaceted roles, 2’-*O*-methylation acts as both a structural and functional regulator of RNA function and fate.

Most cellular 2’-*O*-methylation is thought to be directed by box C/D small nucleolar RNAs (snoRNAs). SnoRNAs have been classified into several distinct groups based on conserved sequence and structural motifs. Box C/D snoRNAs typically contain two (C and C’) (RUGAUGA) and D (and D’) (CUGA) motifs and canonically target ribosomal and small RNAs via base-pairing interactions. Each box C/D snoRNA contains one or two regions, typically 10-15 nucleotides long, known as guide sequences or antisense elements (ASEs). These sequences are located upstream of the conserved D box (and sometimes also upstream of the D’ box, if present). The nucleotide on the target RNA that is destined for methylation is canonically located five nucleotides upstream of the D box (or D’ box) within the region base-paired with the snoRNA’s guide sequence. Box C/D snoRNAs do not function alone. They associate with a conserved set of core proteins (Fibrillarin, NOP56, NOP58, and SNU13 in eukaryotes) to form a small nucleolar ribonucleoprotein (snoRNP) complex. Fibrillarin is the methyltransferase that performs the modification once the snoRNA has guided the complex to the correct site on the target RNA. While most cellular 2’-*O*-methylation is thought to be associated with box C/D snoRNAs, it has been reported that the HIV genome is 2’-*O*-methylated at multiple sites by human FTSJ3, which acts as a methyltransferase in the host cell, but this snoRNA-independent 2’-*O*-methylation pathway remains poorly characterized^18^.

In humans, 137 out of 267 annotated box C/D snoRNAs^29,30^ have no specified target and are therefore classified as orphan snoRNAs. Possible functions of orphan snoRNAs have been recently reviewed^31^. A number of snoRNAs have been shown to target specific sites on mRNA ^17,32^ and even tRNA^33^. Also, some have been reported to be involved in alternative splicing^34^, alternative polyadenylation ^35^ and other noncanonical processes^36–38^.

Greater in-depth exploration of Nm biological functions has been limited due to the lack of robust screening tools to detect Nm sites. Mapping Nm positions has proven challenging for many reasons, including resistance to hydrolysis^5^, the lack of an antibody that can detect Nm and a possibly low stoichiometry of Nm on mRNA. Several high-throughput Nm detection methods have been developed in recent years, including 2’-*O*Me-seq^39^, RiboMeth-seq ^40–42^, RibOxi-seq^43,44^, Nm-seq^14^, Nm-mut-seq^19^ and NJU-seq^21^. RiboMeth-seq has emerged as the only method that consistently detects all known Nm sites in human rRNAs^40–42^. Most importantly, RiboMeth-seq is the only currently available method for the quantitative evaluation of Nm levels at multiple positions, thus providing information on potential Nm variations and providing Nm profiles. However, RiboMeth-seq cannot be used to identify mRNA sites. Recently, Oxford Nanopore Technologies Direct RNA Sequencing (DRS) has been employed by several groups to investigate transcriptome-wide Nm profiles^45,46^. However, Nanopore-based Nm detection is complicated by signal changes that include read mismatches, deletions, and mutations at neighboring positions. As a result, accurate determination of Nm locations using Nanopore DRS often requires machine learning–based computational analysis and/or comparison with snoRNA knockout conditions. We developed RibOxi-seq, which allows the detection of Nm even from small amounts of RNA, including mRNA and non-coding RNA^43,44^. Like Nm-seq, this method includes multiple iterative cycles of oxidation/beta elimination-dephosphorylation reactions to sequentially eliminate unmodified nucleotides from 3’ ends but leave the Nm sites intact. Finally, only RNA fragments with 2’-*O*-methylated 3’ ends can be ligated to linkers for library preparation (**Figure 1A**). Consequently, Nm sites generate a positive signal and not a lack of signal, in contrast to the methods based on the inhibition of reverse transcription (RT) reactions or resistance to hydrolysis. In this study we have further optimized the RibOxi-seq method (now RibOxi-seq2) to make it more sensitive and reproducible.

**Figure 1.**
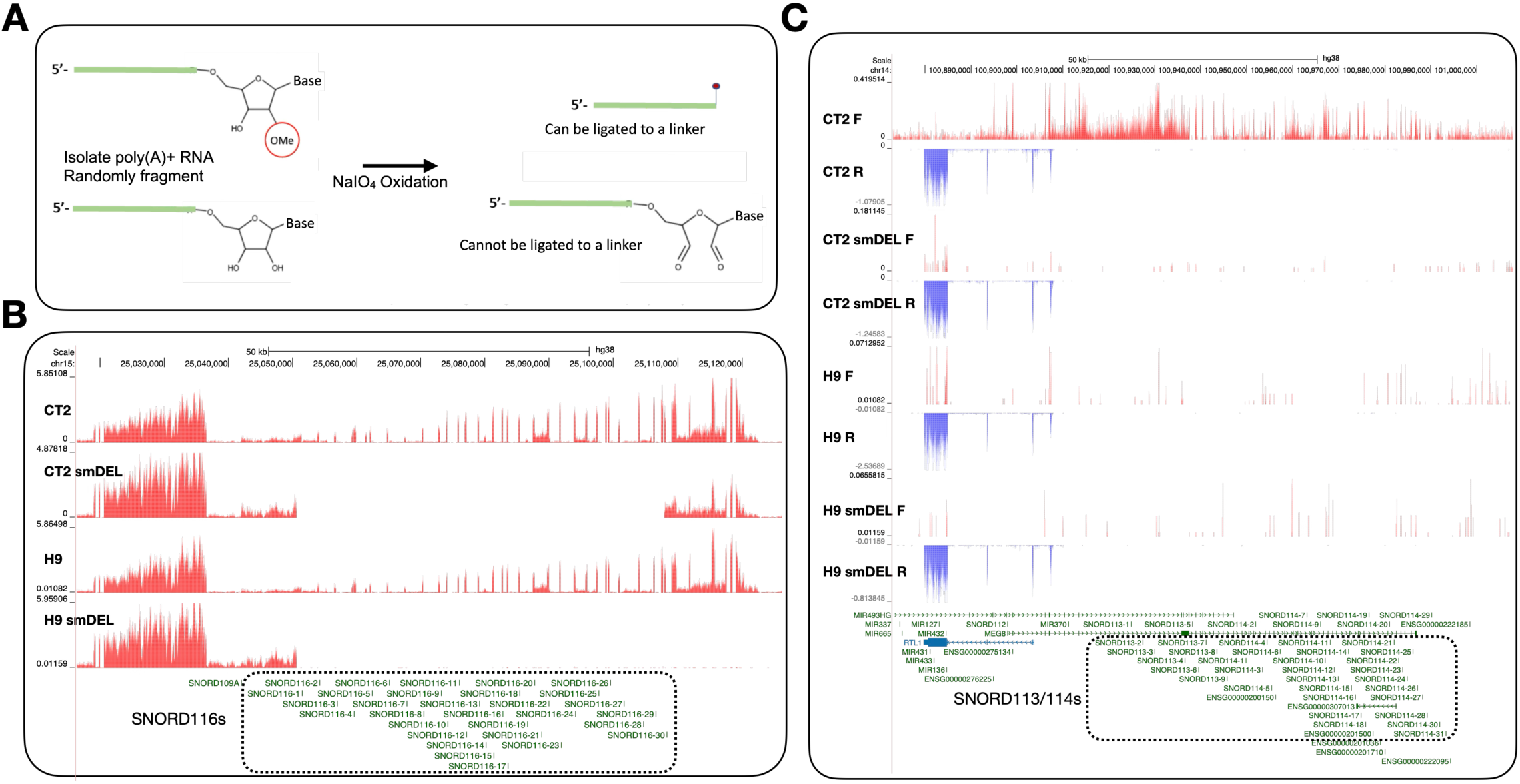
**A.** The RibOxi-seq2 method to map Nm sites. See STAR Methods for details. RNA species with or without a 3′-end Nm modification were subjected to periodate oxidation. RNAs containing a 3′-end Nm modification supported linker ligation, whereas RNAs lacking the modification did not. B. UCSC genome browser image of chromosome 15 illustrating representative bigWig tracks from CT2 and H9 induced neurons and from small-deletion induced neurons. Dashed box outlines the deleted region. **C**. UCSC genome browser image of chromosome 14q32 illustrating representative bigWig tracks from CT2 and H9 induced neurons and from small-deletion induced neurons. Red tracks represent RNA signal on the sense strand, while blue tracks represent RNA signal on the antisense strand. Dashed box outlines the deleted region.

Validating specific Nm positions has also proved challenging. One sensitive method is RTL-P^47^. When reverse transcription (RT) is performed with low concentrations of dNTPs, the RT enzyme has difficulty incorporating nucleotides opposite a 2’-*O*-methylated residue. This leads to premature termination or pausing of cDNA synthesis one nucleotide before the methylated site. However, this method is not reliable for low-abundance RNAs and some structured RNAs^32^. Recently, a site-specific Nm quantification tool, Nm-VAQ, was described^21^. The Nm-VAQ (2’-O-methylation Validation and Absolute Quantification) method can validate the presence and precisely quantify the stoichiometry of Nm at specific RNA nucleotides by leveraging the property that 2’-*O*-methylated RNA is resistant to cleavage by RNase H when it is part of an RNA/DNA hybrid duplex.

Several box C/D snoRNAs have been linked to neurodevelopmental disorders. One of the most notable examples is the orphan C/D box snoRNA cluster SNORD116, which is implicated in Prader–Willi Syndrome (PWS), a rare imprinting disorder affecting approximately 1 in 15,000 newborns^48–50^. PWS results from disruptions of the paternally inherited chromosome 15q11–q13 region, which harbors ∼30 tandemly arranged SNORD116 genes^51–54^. This locus is subject to genomic imprinting, with expression dependent on the paternal allele, and deletions in this region are sufficient to cause the disease. Another imprinted snoRNA cluster with neurological relevance is located on chromosome 14q32.2, where the maternally imprinted SNORD112/113/114 genes locate. Deletions affecting this region give rise to Kagami–Ogata Syndrome, a rare multisystem disorder^55,56^. Intriguingly, SNORD113 and SNORD114 display brain-specific expression patterns, and their strong associations with developmental disorders highlight potential roles in neuronal differentiation and function.

To date, there has been no study published on genome-wide Nm profiling in neurons. To address these knowledge gaps, we have used RibOxi-seq2 to examine isogenic pairs of neurons that only differ in the expression of several orphan box C/D clusters. This allowed us to not only identify neuronal sites of rRNA and small RNA modification but also thousands of mRNA sites, some of which appear to be direct targets of orphan snoRNAs.

## RESULTS

Human embryonic stem cell lines H9 and CT2 were engineered to harbor a doxycycline-inducible neurogenin 2 (NGN2) gene that allows rapid and reproducible differentiation into early postnatal forebrain cortical neurons. Separate isogenic lines were also created in which a small deletion (smDEL) was introduced only on the paternal chromosome 15, removing a cluster of 30 related box C/D snoRNAs (the SNORD116 cluster) that has been implicated in the pathology of Prader-Willi syndrome^50^. The generation of these cells was recently described^57^ and NGN2-induced neurons (iNs) from these cell lines, H9, H9-smDEL, CT2, and CT2-smDEL, have been extensively characterized at the transcriptomic level^58^. Apart from a modest number of overall transcriptomic differences, there are two key features that differentiate the WT and smDEL iNs. First, both the H9-smDEL and CT2-smDEL iNs lack expression of the paternally imprinted SNORD116 cluster of orphan snoRNAs that have been postulated to direct Nms on so far unidentified targets (**Figure 1B and 1C**). Second, although originally derived from parent CT2 cells, the CT2-smDEL cells also differ in their expression of a maternally imprinted locus on chromosome 14q32^59^ and thus differ from CT2 iNs in the expression of the additional orphan SNORD113/114 clusters, comprising 9 and 31 copies, respectively **(Figure 1C**). Importantly, both SNORD113/114 and SNORD116 snoRNAs are expressed in the brain. While the function of SNORD113/114 snoRNAs in the brain is not known, they have been postulated to have a possible role in psychiatric disorders^60^ and deletion of this region leads to the rare disorder Kagami-Ogata syndrome^55,56^. Also, this region has recently been implicated in the maintenance of hematopoietic stem cell self-renewal^61^. Thus, the CT2-smDEL iNs can serve as useful knockout lines for several specific and physiologically relevant subsets of orphan snoRNAs.

### Ribosomal RNA modifications in stem cells and neurons

Given that the relationship between Nm and neuronal disorders remains poorly understood, we sought to gain detailed insights into Nm patterns in an isogenic pair of neuronal populations. RNAs were subjected to RibOxi-seq2 and sites displayed on the UCSC Genome Browser. **Figure 2A-2F** shows representative profiles of 18S and 28S rRNA for CT2, CT2-smDEL, H9, and H9-smDEL iNs. Every peak observed represents an annotated or validated Nm site (**Table S1**). Notably, the peaks are qualitatively consistent between the four cell samples, revealing not only the reproducibility of the RibOxi-seq2 method but also reinforcing the fact that SNORD116 snoRNAs are true orphans as they appear not to target any rRNA Nm. An alternative method, RiboMeth-seq, was next used to corroborate and extend these results (**Figure 2C-2F**). This method has been shown to be highly sensitive and quantitative for rRNA Nm mapping. RiboMeth-seq identified 44 Nm sites in 18S rRNA (3 Nm sites were validated using the site-specific Nm quantification tool, Nm-VAQ^21^) and 67 Nm sites in 28S rRNA, while RibOxi-seq data identified 42 Nm sites in 18S rRNA and 61 Nm sites in 28S rRNA (**Table S1**). In 18S rRNA, using both RibOxi-seq2 and RiboMeth-seq, we identified two Nm sites, Um966 and Um1445, that are known to be highly modified as pseudouridines (Ψ), suggesting the likelihood of double modification (Ψm) at these sites (for this cellular context) ^10^. Likewise, in 28S rRNA, U3818 is known to be modified as Ψm3818, and we also see it as Nm in RibOxi-Seq2 tracks and validated this by Nm-VAQ. One weakness of RibOxi-seq2 is that if there are consecutive Nm sites, only the most 3’ site is observed, and this weakness accounts for most of the discrepancy between the two methods. Thus, 61/63 detectable sites in 28S rRNA were observed by RibOxi-seq2. In order to compare rRNA modifications between pluripotent cells and neurons, we also performed RiboMeth-seq on undifferentiated H9 cells (**Figure 2C and 2D; Table S1**). Nm profiles between H9 cells and H9 iNs were essentially the same except that 18S U354 is unmethylated in stem cells and largely methylated in iNs **(Figure 2C-2F**). These data reveal dynamic ribosomal RNA patterns across different cell lines, suggesting that specific modification levels may contribute to cell-type-specific differentiation. Interestingly, 18S Um354 has previously been associated with differentiation and is directed by SNORD90^62^.

**Figure 2.**
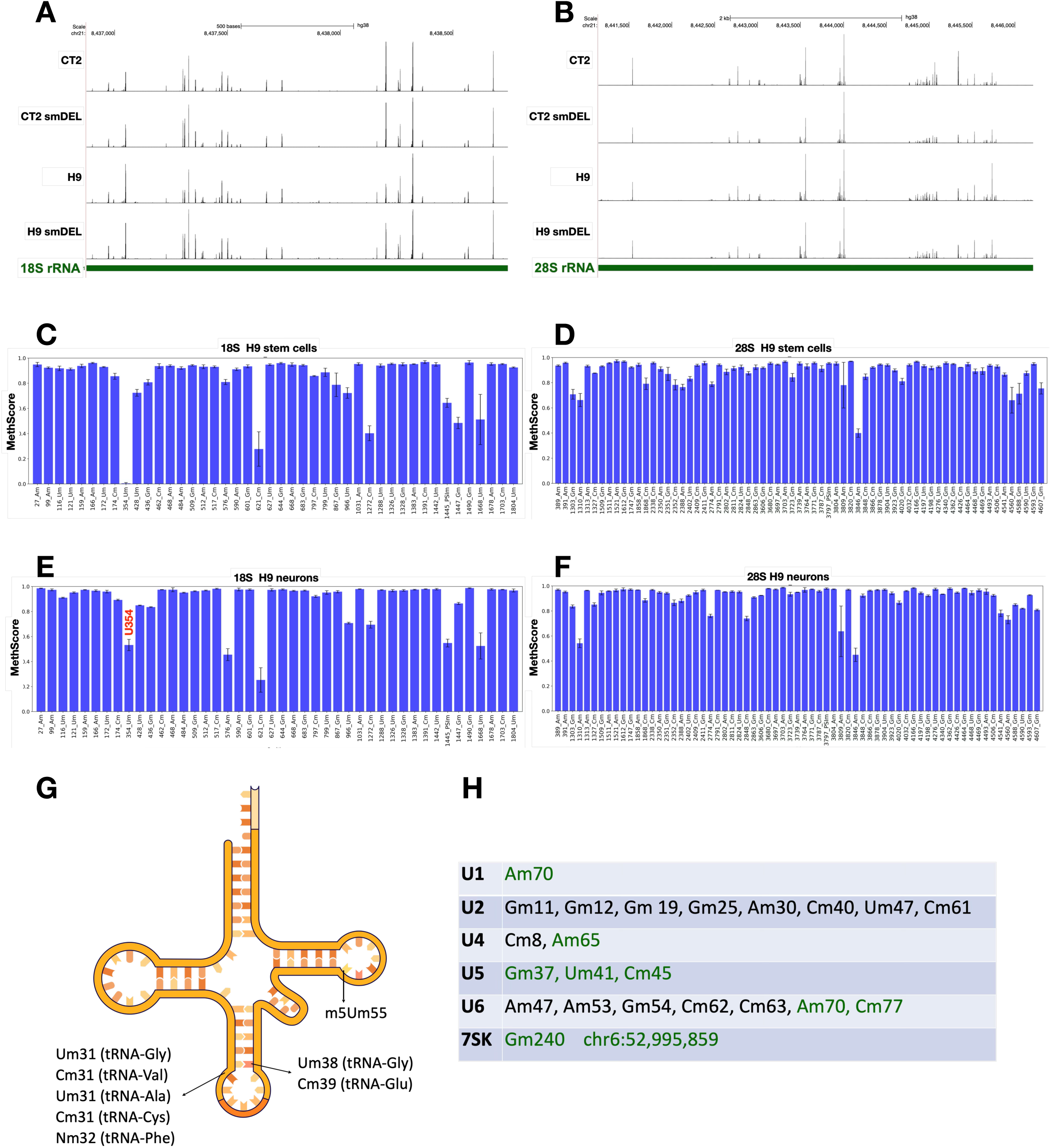
Nm profiling of rRNA. **A**. RibOxi-seq2 peaks in 18S rRNA from induced neurons. Every peak shown aligns with a validated or annotated site. **B**. RibOxi-seq2 peaks in 28S rRNA from induced neurons. Every peak shown aligns with a validated or annotated site. **C**. RiboMeth-seq results for 18S rRNA from H9 HESCs. **D**. RiboMeth-seq results for 28S rRNA from H9 HESCs. **E**. RiboMeth-seq results for 18S rRNA from H9 induced neurons s. Note U354m. **F**. RiboMeth-seq results for 28S rRNA from H9 induced neurons. Positions are numbered; nucleotide identity is indicated. **G**. Nm sites identified in tRNAs. Supporting data is in Figure S2. **H.** Nm sites detected by RibOxi-seq2 in snRNAs and 7SK RNA are highlighted in green, while reference Nm sites are indicated in both black and green. Supporting data is in Figure S2.

It was recently reported that SNORD113/114 snoRNAs may maintain hematopoietic stem cell self-renewal at least in part by altering rRNA Nm levels^61^. Here we observe no 18S or 28S rRNA Nm differences between CT2 and CT2 smDEL iNs, which differ in expression of SNORD113/114. Nm profiles were also analyzed in 5.8S rRNA species and no differences in Nm were detected.

### Small RNA modifications in neurons

RibOxi-seq2 revealed Nm sites in a number of tRNAs. For example, the first nucleotide in the anticodon loop was 2’-O-methylated in tRNA-Gly-CCC-2-2 (Um31), tRNA-Val-CAC-1-4 (Cm31), tRNA-Ala-AGC-2-1 (Um31) and tRNA-Cys-GCA-2-2 (Cm31) (**Figure 2G and S1**). This position is highly conserved compared to the Nm at nucleotide 32 reported in tRNA-Phe, and has been shown to be enzymatically catalyzed by the protein FTSJ1 in human (yeast homolog Trm7), or mediated by SNORD97 or SNORD133 in a snoRNA-dependent fashion^33^. Since these Nm in the anticodon region were not mediated by SNORD113/114/116 snoRNAs, no differences were observed between H9/CT2 wild type and smDEL iNs. Further, Um38 in tRNA-Gly-CCC-2-2 and Cm39 in tRNA-Glu-CTC-1-7 were identified at the first nucleotide following the anticodon loop (**Figure 2G and S1**). These modifications are conserved and correspond to previously annotated (Um/Gm/Ψm39) ^63^. The Nm at position 39 was reported to be catalyzed by a snoRNP complex^64^. As expected, m5Um (2’-O-methyl-5-methyluridine) residues located at the first nucleotide in the T-loop region were also reproducibly captured, including tRNA-Lys-CTT-2-5 (m5Um-54), tRNA-Leu-AAG-2-4 (m5Um-63), tRNA-Gly-GCC-2-4 (m5Um-52), tRNA-Glu-CTC-1-7 (m5Um-53), and tRNA-Asn-GTT-1-1 (m5Um-55) (**Figure 2G and S1**). The m5Um identity and position match previous mass spectrometry results ^63^, indicating that the m5Um modification also resists sodium periodate treatment, and RibOxi-Seq2 cannot clearly differentiate between Nm and m5Um modifications since both are 2’-O-methylated nucleotides.

In addition, reproducible Nm sites were identified in U1 (Am70), U4 (Am65), U5 (Gm37, Um41, Cm45), and U6 (Am70, Cm77) snRNAs (**Figure 2H).** The sites extend our recent work using 2-seq to map Nm sites on U6, supporting the robustness of RibOxi-seq2 in mapping Nm even in small RNA species^65^. Nm modifications in U2 snRNA were not detected, most likely due to the RNA fragmentation procedure failing to enrich small U2 snRNA fragments. Interestingly, we found a novel Nm site in 7SK RNA, Gm240 (**Figure S1**). This site is two nucleotides downstream of A238 which has been reported to be modified as m6A^66^. As no Nm sites have been reported previously in 7SK RNA, we don’t yet know whether this modification occurs in cells other than neurons. Collectively, the RibOxi-seq2 method faithfully identified Nm sites in rRNA, tRNA, and snRNA. As expected, no Nm level changes were found in small RNAs between wild type and smDEL iNs. Together, these results suggest that RibOxi-seq2 may prove useful for mapping many Nm sites even in small RNAs.

### Widespread and robust modifications in mRNAs

A recent study identified thousands of Nm sites in mRNAs of human and mouse cell lines using NJU-seq and validated a number of them using Nm-VAQ^21^. Of thousands of Nm sites identified, about a third were conserved between different cell types. These authors revealed a broad distribution of Nm sites on mRNAs and observed that in their system most validated Nm sites were methylated at ratios from 1% to 30%. Here using RibOxi-seq2 we have also identified thousands of likely Nm sites within mRNAs and lncRNAs (**Table S2**). Importantly, this approach yielded consistent Nm peaks across H9, CT2, and smDEL cell lines, providing strong evidence for the robustness of the method (**Figure 3A**). Owing to a higher sequencing depth in the CT2 samples, we could identify more sites in the CT2 data (989 Nm sites with Nm score 1000 or higher) than in the H9 data (736 Nm sites with Nm score 1000 or higher). When we selected the top 500 sites between both cell types, it was clear that a majority of identified Nm sites are shared between the two parental cell lines (**Figure 3B**). Also, for both H9 and CT2 iNs, Nm profiles were almost identical between the WT and smDEL cells, again highlighting the reproducibility of RibOxi-seq2. Overall, about 40% of Nm sites were in 3’-UTRs, 5% in 5’-UTRs, and the rest in coding exons (**Figure 3C**). While most are shared, some Nm sites were observed only in H9 iNs or CT2 iNs. One example of this is shown in **Figure 3D**. UCHL1 exhibits a strong site in H9 iNs but not in CT2 iNs. Also, the major TOP1 Nm site is in exon 9 in H9 cells but in exon 12 in CT2 cells (**Figure 3E**). Gene ontology analysis revealed that Nm sites are broadly distributed in biological processes, with no clear preference for any pathway (not shown).

**Figure 3.**
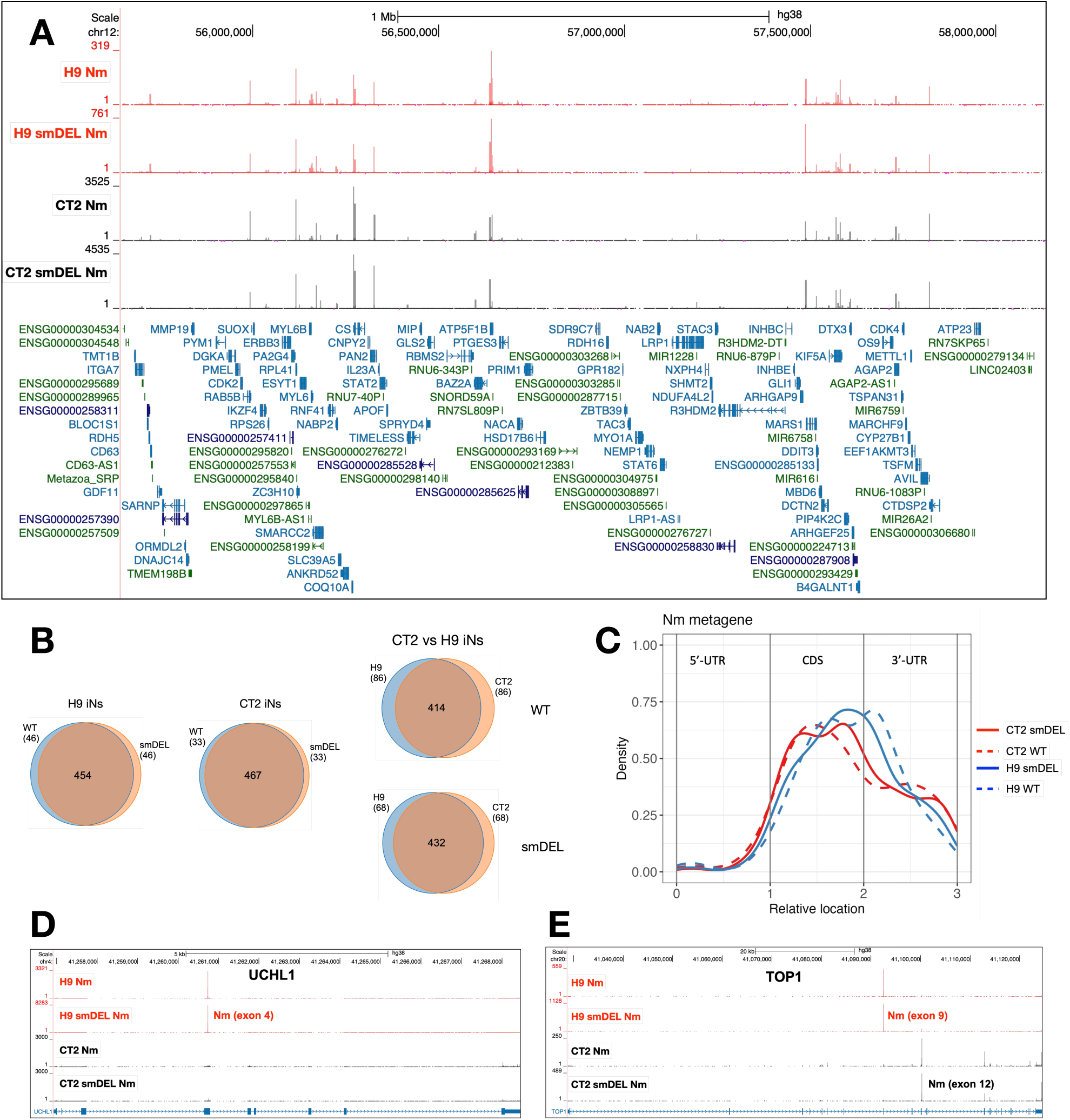
**A**. An example of the reproducibility of RibOxi-seq2. UCSC Genome Browser image of a 2 Mb region on chromosome 12 showing representative bigWig tracks from CT2 and H9 induced neurons and from small-deletion induced neurons. RibOxi-seq2 peaks indicate Nm positions, and gene annotations are displayed below. **B**. Venn diagram illustrating overlapping Nm sites across induced neuron samples; based on the top 500 Nm peaks. **C**. Metagene profile of Nm sites in human mRNA, based on the top 500 Nm sites. **D, E**. Representative examples of two genes with differing Nm positions between H9 and CT2 induced neurons.

Next, we employed Nm-VAQ to validate the modification levels at several mRNA Nm sites. Results are shown in **Figure 4**. Surprisingly, a number of neuronal mRNAs exhibited very high degrees of methylation, in some cases approaching complete modification (**Figure 4F**). For instance, two sites in ACTB exons 4 and 5 were methylated at 97% and 100% (**Figure 4A**), respectively (N.B., RibOxi-seq2 Nm browser peaks as displayed are not quantitative); a site in the 5′-UTR translational control element of TPT1 was modified at 88% (**Figure 4C**); and a site in SNCG exon 3 showed 100% modification (**Figure 4D**). In addition, we detected a site in NUDT21 exon 1 at 50% (**Figure S2**) and another in the 3′-UTR of FAM171A1 at 43% (**Figure 4B**), indicating that not all sites are heavily modified. By contrast, certain Nm sites displayed much lower modification levels. For example, a site in MT-CO1 was modified at only 6.7% and one in FGF13 at 24% (**Figures 4E and S2**). Taken together, these results demonstrate that, at least in iNs, mRNA Nm modifications occur across a wide dynamic range, from nearly absent to fully saturated. Additionally, the Nm sites we identified in neuronal cell lines did not overlap with the mRNA Nm sites previously reported in HeLa cells^21^, suggesting dynamic and/or cell line–specific Nm regulation and variation, or methodologic differences.

**Figure 4.**
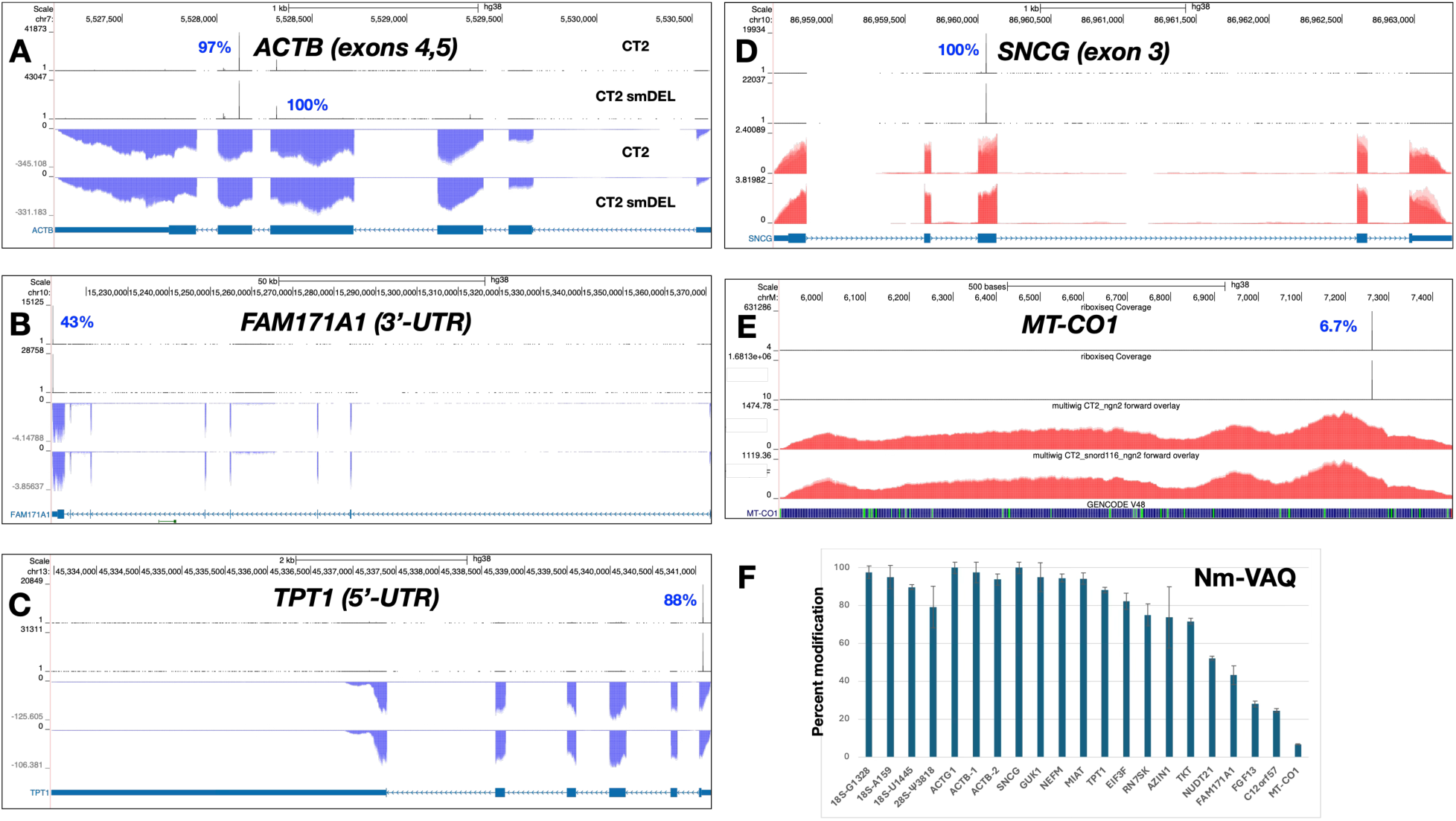
Representative examples of genes displaying differing Nm positions in CT2 and in small-deletion induced neurons. Percentages adjacent to the peaks denote modification levels quantified by Nm-VAQ. Red tracks represent RNA signal on the sense strand, while blue tracks represent RNA signal on the antisense strand. **A**. *ACTB*; Nm peaks in exons 4 and 5. **B**. *SNCG*; Nm peak in exon 3. **C**. *FAM171A1*; Nm peak in 3’-UTR. **D**. Nm peak in *MT-CO1*. **E**. *TPT1*; Nm peak in 5’-UTR. **F**. Quantitation of Nm peaks using Nm-VAQ. Additional data are in Table S3.

#### Lack of canonical snoRNA targeting of most mRNA sites

While all rRNA and most snRNA Nm sites are known to be the targets of box C/D snoRNAs or scaRNAs, this has not been shown yet for most mRNA sites. Our large number of shared Nm sites from our four iN cell lines offered the possibility to test this possibility. We selected CT2 sites with Nm scores above 500 (over 500 sites) and analyzed them by SnoScan^67^ for canonical complementarity to known snoRNAs. Thus, successful hits must exhibit significant complementarity to snoRNAs with the Nm site positioned 5 nucleotides upstream of the snoRNA box D or box D’ (CUGA) motif. Surprisingly, almost none of the abundant shared sites satisfied these criteria. Thus, overall, the vast majority of Nm sites on this list could not be assigned to canonical snoRNA interactions. However, some sites not shared between cells and of lower expression levels could be matched (see below). We next explored the possibility that noncanonical snoRNA-mRNA interactions may play a role in some Nm modifications. While the abundant U3, U8 and U13 snoRNAs have not been previously associated with methyltransferase activity, a recent report studying mRNA-snoRNA physical interactions (sno-KARR-seq) revealed that a high proportion of mRNAs apparently associated with U3 snoRNA, although not necessarily involving canonical basepairing^68^. Further, those authors reported that U3 snoRNA knockdown led to a significant decrease in overall cellular Nm levels. Another group investigating crosslinking of snoRNA-mRNA hybrids also noted a very high number of U3-mRNA interactions^69^. Together, these studies raised the possibility that this snoRNA may have a role in the cell other than in rRNA processing. Interestingly, the sequences around many Nm sites could be aligned with U3, U8 (SNORD118) or U13 snoRNAs by searching for antisense complementarity with snoRNAs catalogues in the snoDB^30^. Comparison of minimum folding energy (MFE) across snoRNAs and RNA regions encompassing each Nm site (±7 nt) (**Figure S3**) revealed a possible connection between U3 snoRNA and many Nm sites.

U3 snoRNA participates in the first stage of rRNA processing^70^ and has been reported to exist in multiple molecular complexes, some containing the canonical box C/D components (Fibrillarin, NOP56, NOP58, and SNU13/15.5K). U3 snoRNA complexes have also been reported to shuttle between the nucleus and cytoplasm^71^. Consistent with a model in which noncanonical snoRNAs directly guide a subset of the Nm sites detected in our RibOxi-seq2 dataset, several of the most prominent Nm peaks overlapped with snoRNA–mRNA chimeras captured by U3 or U8 (SNORD118) in published 293T CLASH data (doi.org/10.64898/2025.12.10.693487). Four representative examples are shown in **Figure 5** and additional overlaps of CLASH fragments with Nm peaks are shown in **Figure S4**. For GUK1, a discrete Nm peak in both H9 and CT2 cells aligns exactly with a U3-derived CLASH fragment, and IntaRNA^72^ predicts a stable U3–GUK1 interaction spanning the modified nucleotide with an interaction energy of −10.5 kcal/mol (**Figure 5A**). A similar pattern is observed at RPL7A, where a strong Nm peak corresponds to a hybrid captured with SNORD118, again supported by a favorable predicted interaction (–9.9 kcal/mol) (**Figure 5B**). Two additional mRNAs, AZIN1 and KMT2A, show the same concordance: each features a sharp Nm peak detected by RibOxi-seq2 that overlaps with U3-captured CLASH reads, and each forms a thermodynamically stable interaction with U3 in IntaRNA modeling (**Figure 5C and 5D**). Notably, for all four transcripts, the strongest predicted base pairing occurs directly over, or immediately adjacent to the Nm-modified position, although these interactions lack the canonical alignment with a recognizable D-box element. Together (**Figures 5 and S4**), these examples are consistent with a model that a subset of mRNAs may acquire 2′-O-methylation through direct interaction with snoRNAs such as U3 and SNORD118, via a noncanonical mechanism.

**Figure 5.**
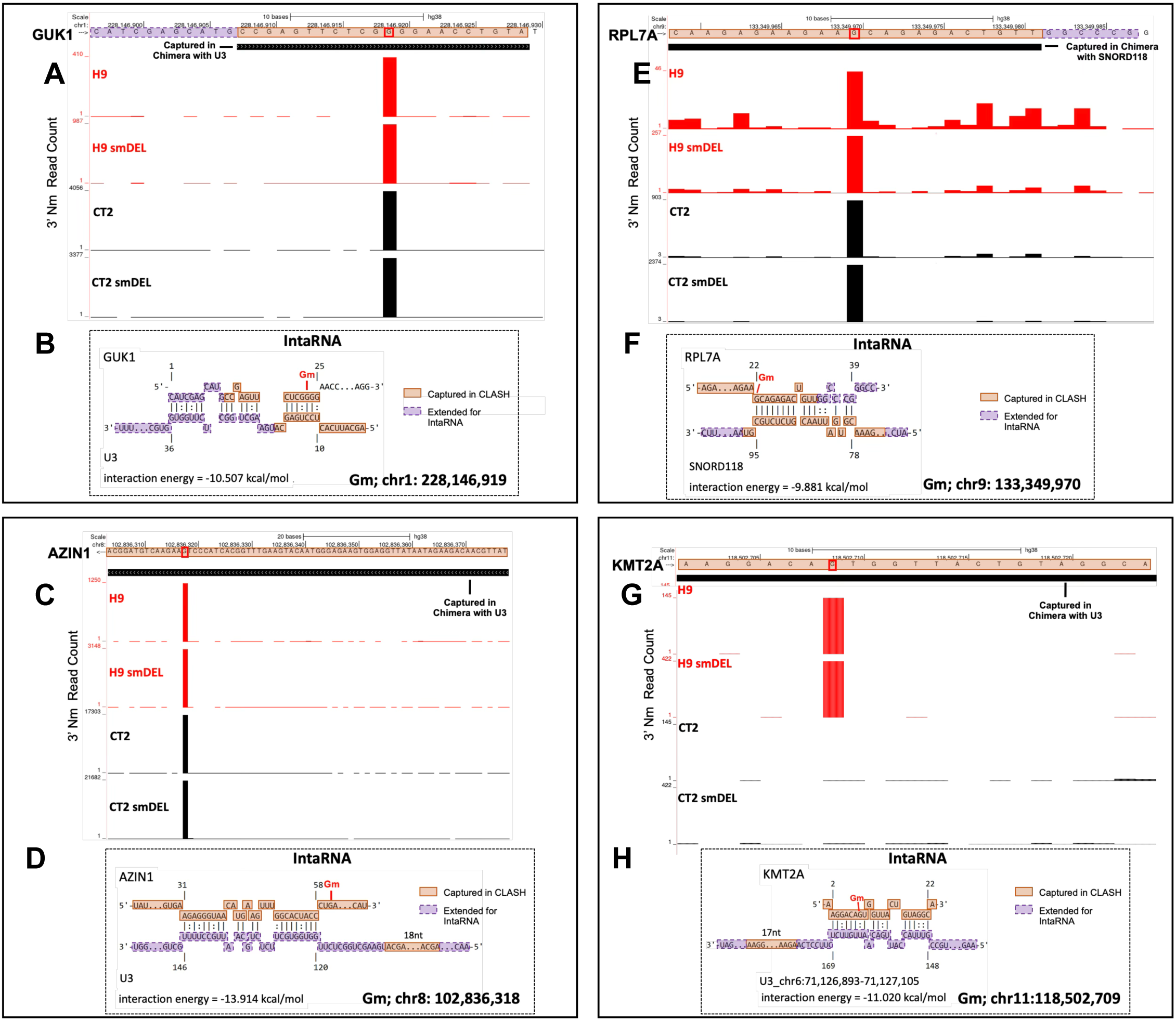
RibOxi-seq2 Nm mapping and snoRNA-guided IntaRNA predictions for sites captured in snoRNA-specific CLASH. (**A, C, E, G**) RibOxi-seq2 tracks show 3′ Nm read counts across four neuron cell line conditions: wild-type (H9, CT2) and SNORD116 deletion lines (H9 smDEL, CT2 smDEL). Black boxes overlaid on the tracks mark RNA fragments captured in chimeras with U3 snoRNA (**A, E, G**) or SNORD118 (**C**). The red signal indicates Nm enrichment peaks detected by RibOxi-seq2. (**B, D, F, H**) Corresponding IntaRNA-predicted base-pairing interactions for the four sites with CLASH overlap. Orange boxes denote regions captured in CLASH hybrids of the target RNA and snoRNA, while purple dashed boxes mark adjacent sequences extended for IntaRNA interaction analysis. Calculated interaction energies (kcal/mol) are shown below each model. Genomic coordinates for the modified nucleoside (Gm) are indicated beneath each panel.

### Targets of orphan SNORD113/114 snoRNAs

Since CT2 iNs and CT2-smDEL iNs differ in expression of the orphan SNORD113/114 snoRNA clusters (**Figure 1B**), we asked whether these cells could be used to identify novel targets for these specific snoRNAs. Our approach was to compare Nm profiles, searching for Nm signals in CT2 iNs that are missing in CT2-smDEL iNs as well as in both H9 iNs and H9-smDEL iNs, which also lack expression of the SNORD113/114 region. This was facilitated by the reproducibility of our data. **Figure 6A** shows a genomic region with a strong peak in the C12orf57 3’-UTR in CT2 iNs that is completely lacking in the CT2-smDEL data. This site is complementary to SNORD114 with the Nm positioned 5 nucleotides upstream of the box D element, consistent with this site being a canonical target. Nm-VAQ revealed that this site is modified in 24% of the transcripts. Another (**Figure 6B**) in the lncRNA MIAT is seen in 94% of transcripts. A similar strategy allowed us to identify additional SNORD113 and SNORD114 targets including PCDH10, SLC22A17, ANKRD13B, SMG5, PARP6, DDX17 and a number of noncoding transcripts (**Figure 6C**).

**Figure 6.**
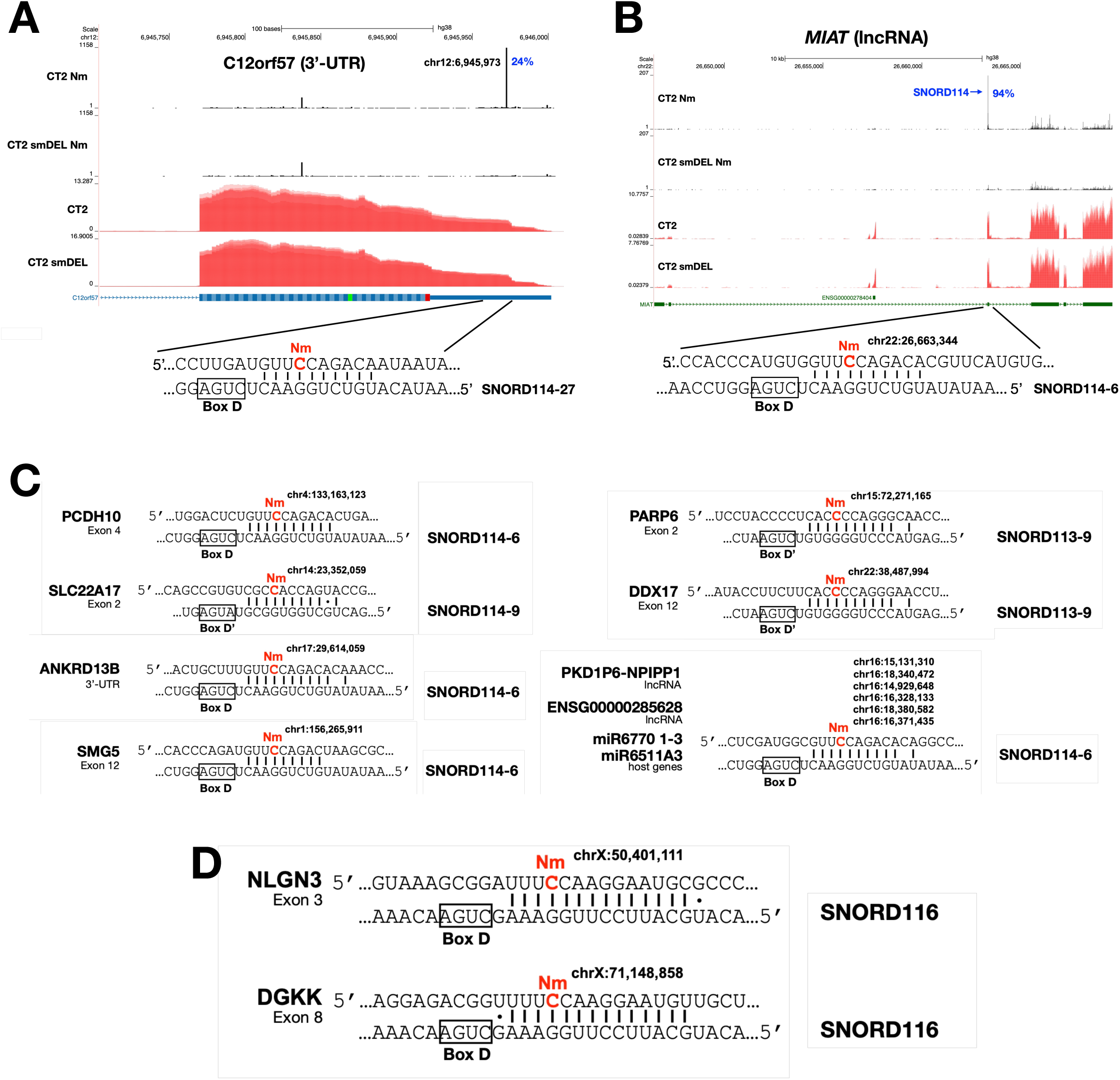
Examples of genes displaying differing Nm positions in CT2 and in small-deletion induced neurons. **A.** *C12ORF57*. Percentages beside the peaks denote modification levels quantified by Nm-VAQ, and the corresponding gene expression profiles are shown in red. **B**. *MIAT*. Percentages beside the peaks denote modification levels quantified by Nm-VAQ, and the corresponding gene expression profiles are shown in red. **C.** Additional SNORD113/114 targets identified. Base pairing between mRNAs containing Nm sites (highlighted in red) and their respective snoRNAs (SNORD114 and SNORD113). Canonical C/D box sequence motifs are boxed. **D.** SNORD116 targets identified.

### Targets of orphan SNORD116 snoRNAs

Since lack of expression of SNORD116 snoRNAs has been related to Prader-Willi syndrome, the smDEL cell lines with SNORD116 cluster deletions represent a valuable resource to investigate the molecular basis of the disease. In order to qualify as potential canonical sites, there needed to be an Nm signal in both H9 and CT2 iNs but no signal at the same position in both H9-smDEL and CT2-smDEL iNs. Also, the Nm sites needed to be complementary to SNORD116 with the Nm site positioned 5 nucleotides upstream of a box D or D’ element. While we found a number of SNORD113/114 targets, we were only able to identify two potential SNORD116 targets. However, the two targets identified are of considerable interest. These are NLGN3 and DGKK (**Figure 6D**). For each of these mRNAs, there is extensive basepairing with multiple SNORD116 species and the potential Nm sites are 5 nucleotides upstream of the box D element. However, as PWS pathology is closely connected to lack of expression of SNORD116s, our failure to identify numerous robust SNORD116 targets was unexpected. This could be for several reasons. First, some physiological SNORD116 targets may not be expressed well in our iNs. Since PWS is generally thought to be primarily a disorder affecting the hypothalamus, some targets most abundantly expressed there could be poorly expressed in iNs. Second, our findings could indicate that these snoRNAs, which are highly abundant in neurons, may have additional important functions apart from guiding 2’-O-methylation. Third, some PWS pathology may be connected not directly to SNORD116 snoRNAs, but to expression of other noncoding transcripts expressed from this genomic region^73–75^.

## DISCUSSION

Here we have used RibOxi-seq2 to generate the first comprehensive picture of the Nm landscape in neurons. Sequencing was carried out on two isogenic pairs of NGN2-induced neuronal cultures derived from embryonic stem cell lines. Each isogenic pair included one wild-type cell line and another harboring a small deletion of 30 related tandem orphan snoRNAs (SNORD116s) from the paternal chr15q11-q13 region, whose loss of expression is associated with Prader-Willi syndrome. The CT2-based isogenic pair also differed in expression of the orphan SNORD113/114 clusters from the imprinted chr14q32.2 region. Thousands of shared Nm sites were identified in these cells, in both coding and noncoding RNAs. The most abundant cellular Nm modifications occur in rRNA. In rRNA, RibOxi-seq2 identified almost all annotated Nm sites but is limited by two technical issues. First, when two Nm sites exist in tandem, RibOxi-seq2 can only see the distal site, except when this site is not fully modified. This led to an inability to monitor several sites in both 18S and 28S rRNA. Second, since 2-seq2 is qualitative and not quantitative, sites modified at a low level even in abundant RNAs such as rRNA can lead to some Nm peaks that are not dramatically higher than background signals. Owing to these issues, we further analyzed ribosomal Nm profiles using RiboMeth-seq, which does not suffer from these drawbacks. Results showed not only that RibOxi-seq2 is highly efficient at identifying rRNA Nm sites, but also that ribosomal RNA in iNs has a distinct modification signature compared to that in embryonic stem cells. In particular, Um354 in neuronal 18S rRNA was seen by both methods and has been reported to be more highly expressed in differentiated cells and directed by SNORD90^62^. This raises the question of how a specific rRNA Nm profile affects translational efficiency or regulation in neurons. Might it affect the translation not only of neuronal mRNAs, but also those harboring neuron-specific Nms? Apart from rRNA, we also observed a number of reported Nm sites in other abundant noncoding RNAs such as snRNAs and tRNAs, showing that RibOxi-seq has the potential to monitor such sites in RNA samples. We also identified and validated a novel Nm site in 7SK RNA.

Owing to its ability to detect Nm sites even in low-abundance RNAs, RibOxi-seq2 identified thousands of Nm sites in hundreds of mRNAs and lncRNAs. Importantly, a large fraction of these sites appeared in all four iN cultures, while only a small fraction of Nm sites appeared to be cell- or parent-specific. We don’t yet know the basis of specificity, but we hypothesize that the shared Nm sites include a neuronal signature of genome-wide 2’-O-methylation. While most Nms are shared between the different neuron samples, some are cell-specific. Nm sites in mRNAs are broadly distributed both in functional categories and in regions of internal RNA modification. Thus, about 5% are in 5’-UTRs, 40% in 3’-UTRs, and the rest in coding exons. While some mRNAs may harbor sites that are partially modified, other mRNAs have Nm sites that are almost fully modified. Nm-VAQ revealed that a number of sites in abundant mRNAs are present at almost 100% modification. Such high levels of modification have also been reported recently by others^20^, though most sites identified in this study do not match the positions reported in that work and may thus represent important cell-specific RNA modifications or differences in methodology.

What targets most mRNA Nm sites? In our studies, we were surprised to discover that we could not match the great majority of mRNA Nm sites to canonical RNA-snoRNA interactions. This suggested that such modifications may occur via noncanonical interactions, or that many may be directed via recruitment of other activities such as the methyltransferase FTSJ3. Intriguingly, however, we observed that many sites exhibit complementarity to abundant but noncanonical snoRNAs such as U3, U13, and U8 (SNORD118). Do these snoRNAs and their associated proteins harbor methyltransferase activities or have the ability to recruit them? U3 has been reported to exist in multiple distinct snoRNP complexes, and U3 as well as some of its protein components have been shown to shuttle between the nucleus and cytoplasm^71,76,77^. Our findings with U3 are consistent with the recent report of abundant U3-mRNA interactions using the sno-KARR-seq RNA-RNA interaction method^68^. Importantly, those authors observed that knockdown of U3 led to a reduction in overall levels of mRNA Nm modification. Another group also reported a very high number of U3-mRNA interactions^69^. Together with our CLASH results, these findings lead us to hypothesize that U3, and perhaps also U13 and U8, play an important role in mRNA Nm modification in addition to its role in rRNA maturation.

While Nm sites in abundant mRNAs could not be assigned to snoRNA interactions following established rules, our results support the assertion that some Nm peaks do represent canonical snoRNA-mRNA interactions. This possibility is supported by the identification of a number of sites that appear to be canonical targets of previously annotated orphan snoRNAs, including members of the SNORD113, 114, and 116 families. Our data represent the first identification of likely targets of these snoRNAs. Most of these targets are in RNAs that are connected to neurodevelopmental disorders.

Deletion of the SNORD113/114 region (chr14q32.2) leads to the rare multi-symptom disorder Kagami-Ogata syndrome^55,56^. Specific expression patterns of SNORD113/114 snoRNAs in the brain and strong associations with neurodevelopmental disorders point to crucial and specialized roles in brain development, function, and potentially in the pathogenesis of neurological conditions. SNORD113-6 has been reported to methylate tRNA^78^ but our data suggest a possible additional role in mRNA modification. Novel mRNA targets of SNORD113 that we have identified here include PARP6 and DDX17 (**Figure 6C**). PARP6 is a regulator of hippocampal dendritic morphogenesis^79^. DDX17 interacts with and co-regulates the transcriptional repressor element 1-silencing transcription factor (REST), which is a critical regulator of neuronal differentiation, suppressing neuronal gene expression in progenitor cells^80^.

SNORD114 targets also shed light on potential roles in neuronal development. C12orf57 has been reported to be a critical gene involved in brain development, particularly the formation of the corpus callosum and the regulation of synaptic function. It is also involved in neurodevelopment and is a regulator of synaptic AMPA currents and excitatory neuronal homeostasis^81,82^. Mutations in this gene are associated with developmental brain abnormality^83^. MIAT is a lncRNA associated with neuronal development^84,85^. PCHD10, a cadherin-related neuronal receptor, is associated with autism and neuropsychiatric disorders^86,87^. SLC22A17 (Solute Carrier Family 22 Member 17), also known as the lipocalin-2 receptor, has emerged as a significant player in the brain, particularly in the context of neuroinflammation and neurodegeneration. While initially identified for its role in organic cation transport and iron homeostasis, recent research has highlighted its critical involvement in the blood-brain barrier (BBB) integrity and cell death pathways, especially after stroke^88^. ANKRD13B is a ubiquitin-binding protein that is widely expressed in the brain. Finally, it is noteworthy that SNORD114 appears to target a number of noncoding transcripts, including lncRNAs and miRNA host genes. PKD1P6-NPIPP1 is a cell-cycle related lncRNA^89^. Further studies are warranted to investigate the functional ramifications of these Nm modifications.

We also found two potential SNORD116 targets, DGKK and NLGN3, and these genes have functions consistent with roles in PWS. NLGN3 (neuroligin 3) is a crucial postsynaptic cell adhesion molecule that plays a pivotal role in organizing and stabilizing synapses and which is linked to autism^90^. It is also a key regulator of gonadotropin-releasing hormone deficiency and is connected to hypogonadism^90^. DGKK (diacylglycerol kinase kappa) converts diacylglycerol to phosphatidic acid, regulating the balance between these two important signaling lipids in the brain. These lipids are involved in neurotransmitter signaling, synaptic plasticity, and overall neuronal communication^91,92^. DGKK has been reported to be a (and possibly the) primary target of the Fragile X-related protein FMRP^91,93^. Interestingly, both NLGN3 and DGKK are translationally upregulated by FMRP^91,94^ so the translational consequences of Nm modifications in their mRNAs clearly warrant future study. Thus, both of the potential SNORD116 targets identified here may contribute to the developmental, neurological, and behavioral features of PWS. In fact, among others, both of these genes were hypothesized as possible PWS targets in previous work^95^.

We do not yet know the function(s) of Nms in targeted mRNAs. At this time, our data do not allow us to determine any strong correlation between Nms and RNA stability, or any apparent alteration in alternative polyadenylation, as suggested by Li et al. ^20^. We do note, however, that the orphan SNORD113 and SNORD114 targets identified here do not show significant expression differences in iNs that do or do not express these snoRNAs. For example, in cells that do and do not express SNORD114s (CT2 and CT2-smDEL iNs), we see essentially identical expression levels of C12orf57 and MIAT (for example, see expression levels listed for RNA-seq tracks in **Figure 6A and 6B**), although these RNAs differ in modification between the cells. It has been shown that Nms can induce ribosome pausing during translation elongation^16^. While this has the potential to reduce protein expression, it is possible that such pausing, at least at some Nms within mRNA coding regions, may serve another purpose, to promote or facilitate specific protein folding of nascent peptides^96^. Further, Nms within coding or noncoding regions may serve other functions. They may serve to facilitate or inhibit the assembly of protein complexes involved in RNA processing, transport or decay. This could be via stabilization of altered RNA conformations^13^. Finally, the possibility exists that Nms in 3’-UTR elements may interfere with mRNA fate regulation by microRNAs, perhaps by preventing RNA cleavage.

In conclusion, we note that the isogenic cell models described here may prove useful in additional studies of Nm modifications and functions. As these cells are embryonic stem cells, they can be differentiated into a multitude of additional cell and organ types, and their differences in orphan snoRNA expression can be exploited to examine the roles of SNORD113, SNORD114, and SNORD116 box C/D snoRNAs in multiple tissue types, both at transcriptomic and targeted Nm levels.

### Limitations of the study

1. While RibOxi-seq2 identifies many Nm sites, these are mostly seen in highly or moderately expressed transcripts. Sites in mRNAs of low abundance may be less reliably seen.
2. Many PWS targets may remain unidentified in this model, and we may have missed many sites present in other neuronal populations. Also, PWS is thought to be largely a disorder connected to the hypothalamus. This limitation also applies to SNORD113/114 targets, which may exist in numerous other cell types. By differentiating into different cell and organ types, the CT2/CT2 smDEL isogenic cell pair of human ESCs used here may offer the potential to gain even further insights into the functions of SNORD113, SNORD114, and SNORD116. For example, differentiating into hypothalamic organoids may offer the chance to identify additional physiologically relevant SNORD116 targets.
3. A mechanistic connection of U3/U8 (SNORD118)/U13 to Nm is not conclusively proven at this point.
4. We don’t yet know the biological function of Nm modifications and cannot connect Nm to stability, translation, or subcellular localization. For example, we see a strong Nm site in the 5’-UTR of TPT1, positioned within a structured translation regulation TOP element. Could this affect neuronal translation regulation in neurons or other cells?
5. Functional consequences of the neuronal rRNA Nm signature we observe have not yet been elucidated.
6. We don’t yet know when modifications occur (co-transcriptionally or post-transcriptionally) or primarily in the nucleus or in the cytoplasm. This issue is particularly relevant for U3, which has been reported to be at least partially localized to the cytoplasm.
7. Both H9 and CT2 cells are female, and future work should repeat the studies in a male line such as H1.
8. Using the snoDB search tool^29,30^, we noted that a number of Nm sites also appear to show strong complementarity to box H/ACA snoRNAs. This raises the possibility that some Nm sites may share modification by pseudouridylation.

## RESOURCE AVAILABILITY

### Lead contact

Further information and requests for resources and reagents should be directed to and will be fulfilled by the lead contact, Gordon G. Carmichael.

### Materials availability

This study did not generate new unique reagents.

### Data and code availability

RibOxi-Seq2 data for iNs have been deposited at GEO with Bioproject accession number PRJNA1348454. RibOxi-seq data for 293T cells have been deposited at GEO with accession number GSE188194. The scripts for our RibOxi-Seq2 pipeline are available from https://github.com/yz201906/RibOxi-seq. Any additional information required to reanalyze the data reported in this paper is available from the lead contact upon request.

## ACKNOWLEDGEMENTS

This work was supported by NIH grants R35GM118140 to B.R.G., HD099975 and a grant from the Foundation for Prader Willi Research to G.G.C., R35GM146883 to J.D.B, F31HD114435 to S.A.A. and R01GM135383 to C.L.H. We thank Sara Olson for helpful suggestions regarding the Illumina sequencing.

## AUTHOR CONTRIBUTIONS

Conceptualization, G.G.C.; formal analysis, X.Y., Y.L., Y.Z., S.A., B.A.E., and G.G.C; investigation, X.Y., Y.L., Y.Z., S.A.A., B.A.E., and G.G.C; resources, Y.L., Y.Z., S.A. and B.A.E.; writing – original draft, X.Y. and G.G.C.; writing – review & editing, all authors; visualization, X.Y., S.A.A., B.E., and G.G.C.; supervision, C.L.H., J.B., B.R.G. and G.G.C.; funding acquisition, C.L.H., J.B., B.R.G. and G.G.C.

## DECLARATION OF GENERATIVE AI AND AI-ASSISTED TECHNOLOGIES

During the preparation of this work, the author(s) used ChatGPT and Gemini to improve readability. After using this tool or service, the authors reviewed and edited the content as needed and take full responsibility for the content of the publication.

## STAR★METHODS

### KEY RESOURCES TABLE

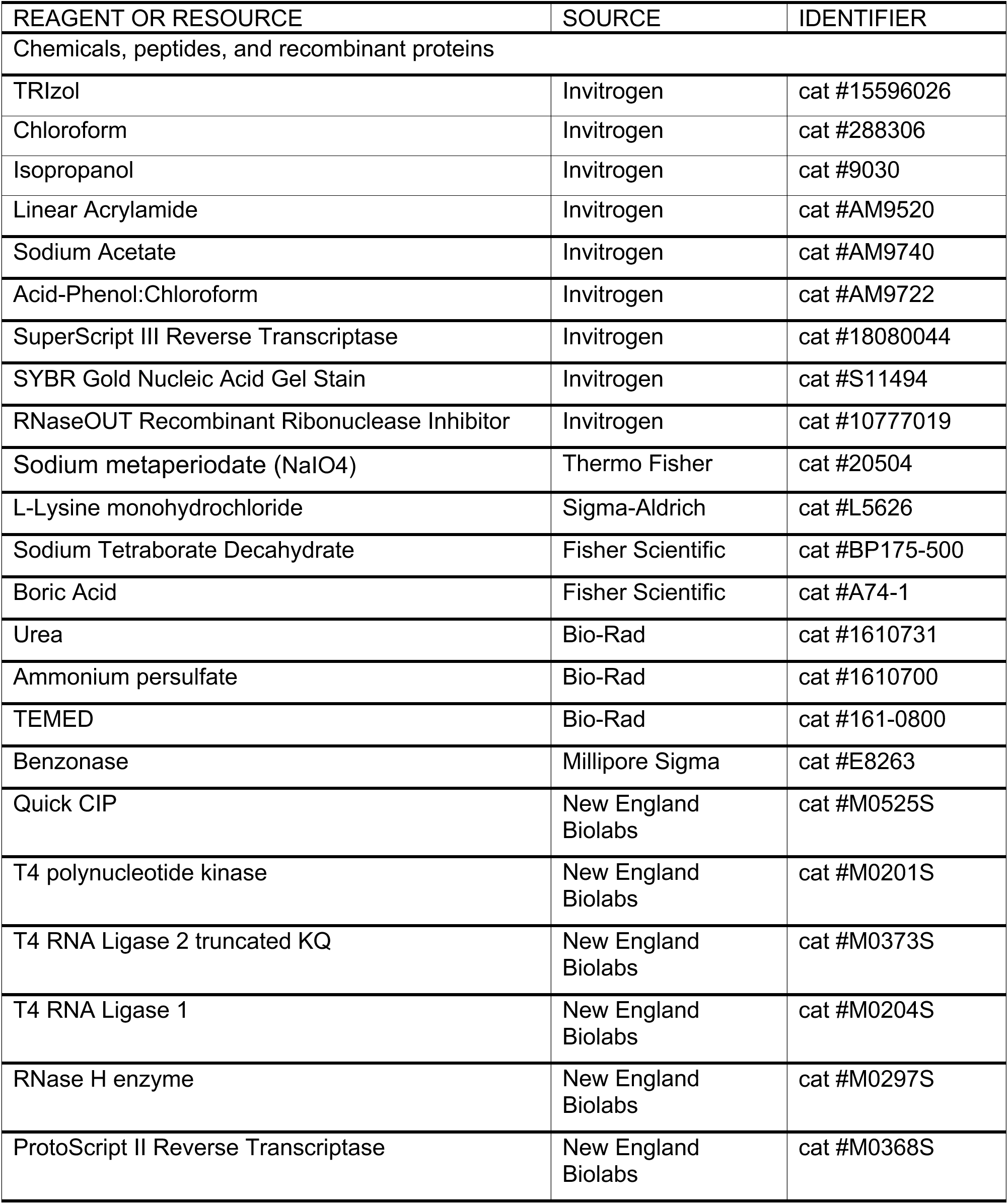
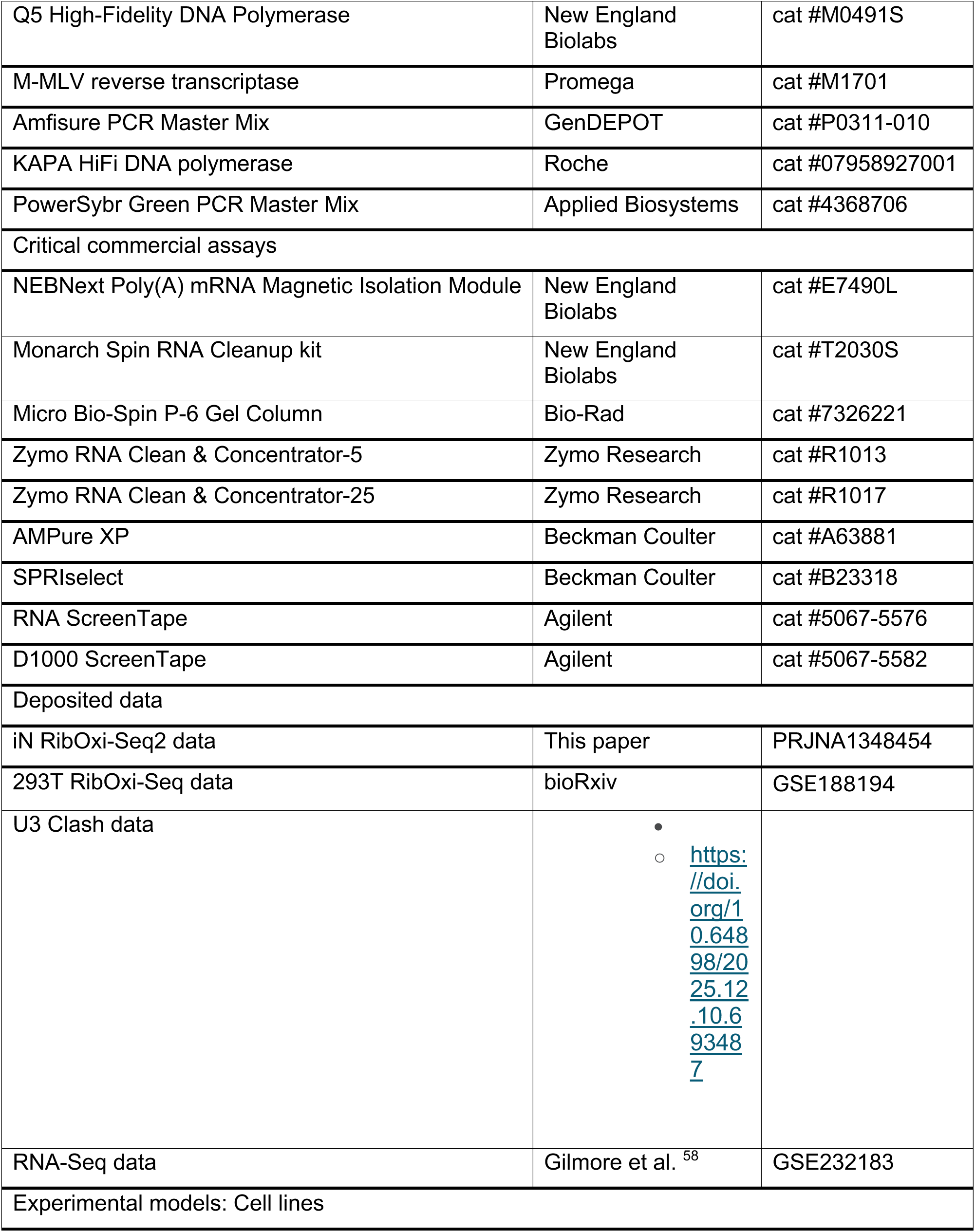
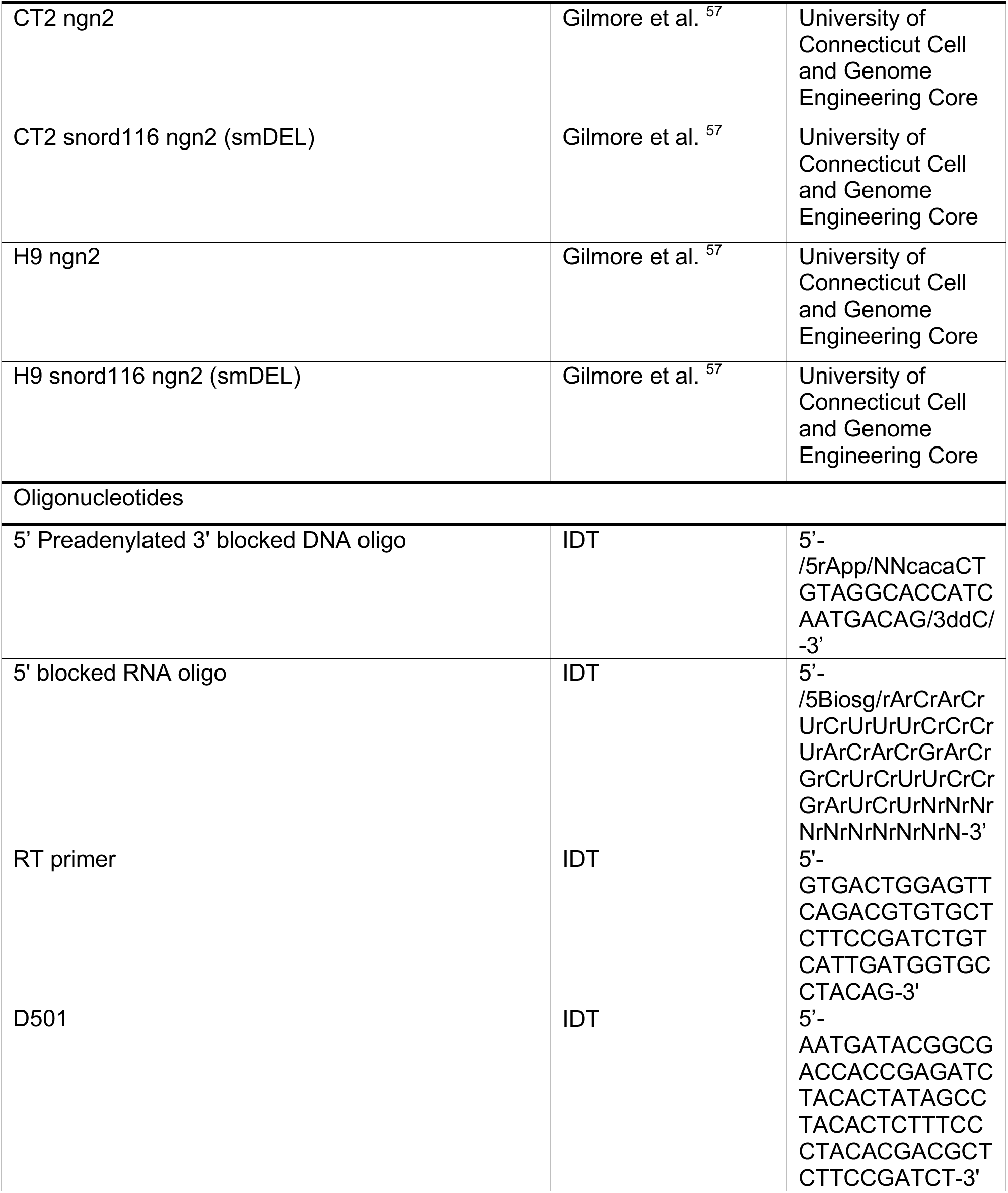
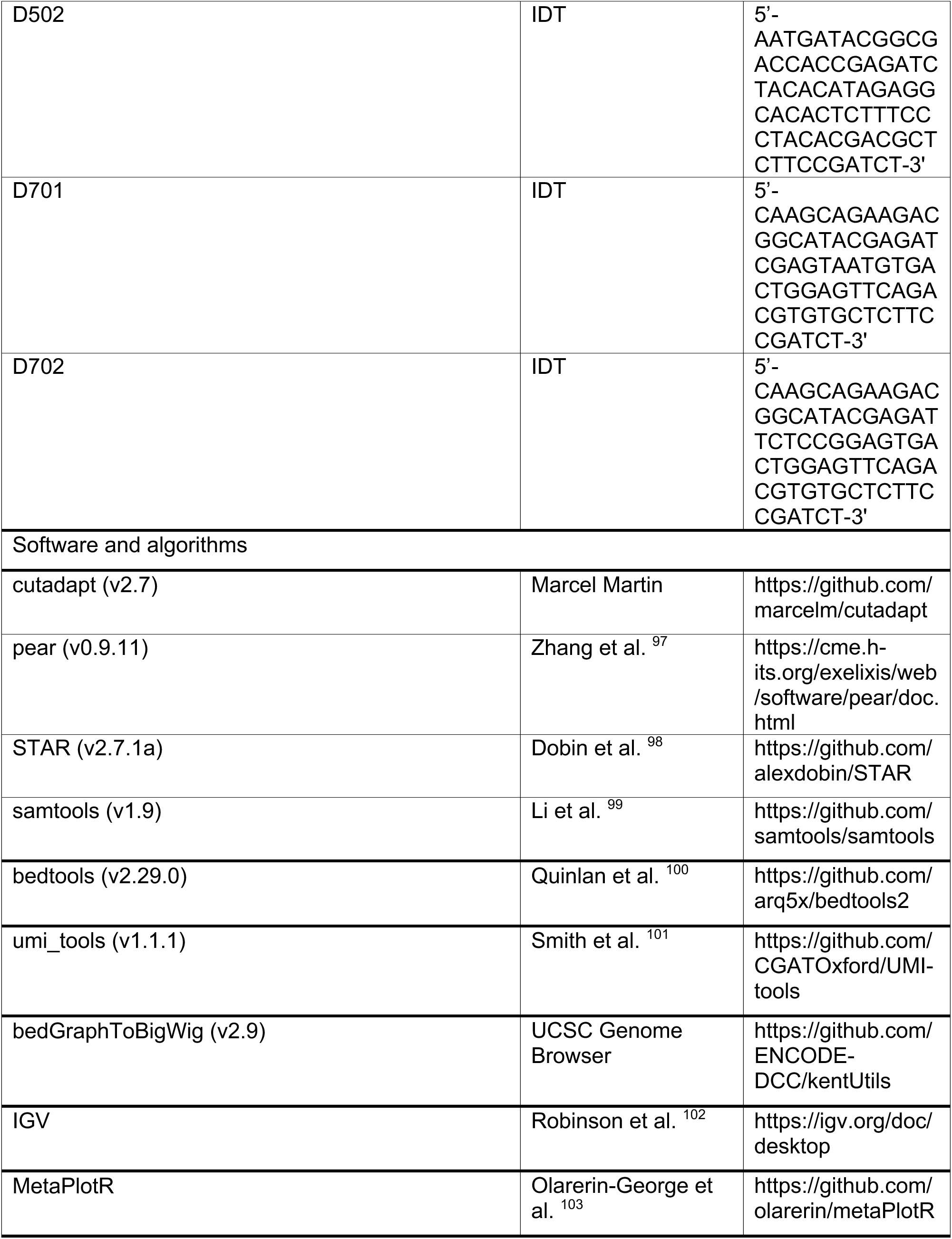
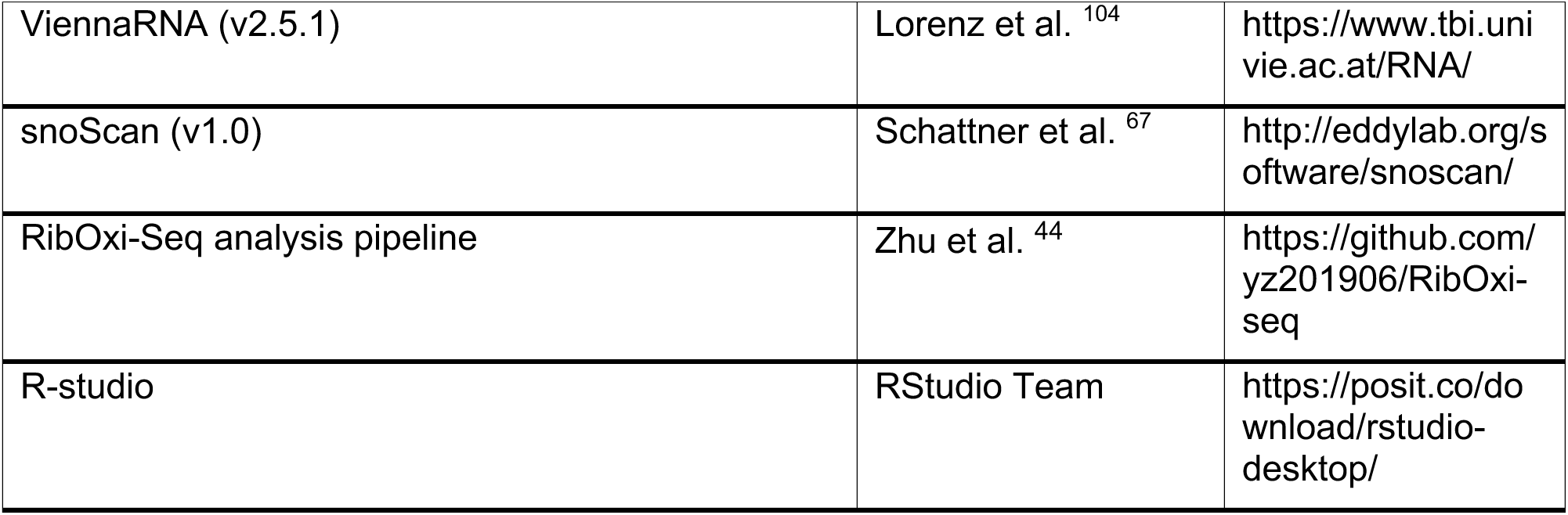

### EXPERIMENTAL MODEL AND STUDY PARTICIPANT DETAILS CELL CULTURE

Culture and differentiation were carried out as described previously^58^. CT2 and H9 human embryonic stem cells (hESCs) including SNORD116 deletion variants were cultured under feeder-free conditions using Matrigel-coated (Corning, #354277) 100-mm dishes and maintained in mTeSR Plus (STEMCELL Technologies, #100-0276) in a humidified atmosphere at 37°C with 5% CO₂. The culture medium was replaced daily, and cells were passaged approximately every 4–5 days upon reaching 80–90% confluency. For passaging, the medium was removed, and dishes were treated with 0.5 mM EDTA (Invitrogen, #15575020) in PBS, followed by a 2-minute incubation at 37°C. The EDTA solution was then aspirated, and fresh mTeSR Plus medium was added. The cell aggregates were carefully detached using either a StemPro EZPassage Tool (Gibco, #23181010) or by gently scratching the dish with a glass pipette. The passage ratio was maintained at approximately 7–10 to ensure optimal growth.

When the hESCs reached 70–80% confluency, cells were prepared for differentiation. First, any differentiated cells were manually removed, and the 100-mm dish was rinsed with PBS. The cells were then treated with Accutase (Millipore, #SCR005) and incubated for 2 minutes at 37°C. After incubation, 10 mL of basal media was added to the cell suspension, which was transferred to a 50-mL conical tube and centrifuged at 1200 rpm for 3 minutes. The supernatant was aspirated, and the pellet was resuspended in 1 mL of Induction Media (IM). The cells were singularized by pipetting up and down 3–5 times using a 1-mL pipette. An additional 9 mL of IM was then added to the suspension.

The Induction Medium (IM) was prepared by supplementing DMEM/F12 (with HEPES) (Gibco, #11330032) with 1X N-2 Supplement (Gibco, #17502048), 1X MEM Non-Essential Amino Acids (Gibco, #11140050), and 1X GlutaMAX Supplement (Gibco, #35050061).

For differentiation, 4 million cells were plated in a Matrigel-coated (Corning, #354277) 100-mm dish and cultured in IM supplemented with 2 μM Doxycycline Hydrochloride (Fisher Scientific, #BP26535). The cells were fed daily with IM containing 2 μM doxycycline hydrochloride for four days.

On day 4 of differentiation, the cells were singularized again using Accutase as described above. The cell pellet was resuspended in Cortical Media (CM), which was prepared by mixing equal volumes of DMEM/F12 with HEPES and Neurobasal Medium (Gibco, #21103049) and supplementing with 1X B27 Supplement (Gibco, #17504044), 10 ng/mL Recombinant Human BDNF Protein (R&D Systems, #248-BD), 10 ng/mL Recombinant Human GDNF Protein (R&D Systems, #212-GD), 10 ng/mL Human NT-3 Recombinant Protein (PeproTech, #450-03), and 1 μg/mL Laminin Mouse Protein (Gibco, #23017015).

Cells were plated at a density of 13 million cells per 100-mm dish in CM supplemented with 10 μM ROCK inhibitor Y-27632 dihydrochloride (Tocris, #1254). A complete medium change with CM was performed the following day, and the media was subsequently changed daily until collection on day 11.

The 100-mm dishes were pre-coated with 100 μg/mL Poly-D-lysine hydrobromide (PDL, Millipore Sigma, #P0899) and 5 μg/mL laminin. Initially, 6 mL of the 100 μg/mL PDL working solution was applied to each dish and incubated overnight to ensure adequate coating. The next day, the dishes were thoroughly washed twice with PBS to ensure they remained hydrated. Subsequently, 6 mL of the 5 μg/mL laminin working solution was added to the PDL-coated dishes, which were then incubated overnight to prepare them for neuron differentiation.

### METHOD DETAILS

#### RNA Extraction

Total RNA was extracted from cells using the TRIzol reagent protocol. Briefly, cells were washed with PBS, pelleted by centrifugation, and lysed in TRIzol reagent (Invitrogen, #15596026). After incubation at room temperature for 5 minutes, 0.2 mL of chloroform (Invitrogen, #288306) per mL of TRIzol was added, followed by vigorous shaking and centrifugation at 12,000 × g for 15 minutes at 4°C. The aqueous phase containing RNA was collected, mixed with 0.5 mL of isopropanol (Invitrogen, # I9030) per mL of TRIzol, incubated for 10 minutes, and centrifuged at 12,000 × g for 10 minutes at 4°C. The resulting RNA pellet was washed with 75% ethanol, air-dried, and resuspended in RNase-free water. RNA cleanup and DNase I treatment were performed using the RNA Clean & Concentrator-25 (Zymo Research, #R1017) according to the manufacturer’s instructions.

#### RNA Ethanol Precipitation

To 100 µL of RNA solution, 1 µL of Linear Acrylamide (250–500X; Invitrogen, AM9520) and 250 µL of ice-cold 100% ethanol (2.5–3.0 volumes) were added. The mixture was thoroughly mixed and stored at −80°C for 1 hour or overnight at −20°C to allow RNA precipitation. The precipitated RNA was recovered by centrifugation at 14,000 × g for 20 minutes at 4°C. The supernatant was removed, and the RNA pellet was washed with 0.5 mL of ice-cold 75% ethanol, followed by centrifugation at 10,000 × g for 5 minutes at 4°C. Residual ethanol was carefully removed using a 20-µL pipette after a brief spin-down, and the pellet was air-dried on ice for 5–10 minutes before being resuspended in nuclease-free water.

#### DNA Ethanol Precipitation

To 100 µL of DNA solution, 10 µL of 3M Sodium Acetate (Invitrogen, #AM9740), 1 µL of GlycoBlue Coprecipitant (Invitrogen, #AM9515), and 250 µL of ice-cold 100% ethanol (2.5–3.0 volumes) were added. The solution was thoroughly mixed and stored overnight at −20°C to allow DNA precipitation. The DNA was recovered by centrifugation at 14,000 × g for 20 minutes at 4°C. The supernatant was carefully removed, and the DNA pellet was washed with 0.5 mL of ice-cold 75% ethanol, followed by centrifugation at 10,000 × g for 5 minutes at 4°C. Residual ethanol was carefully removed using a 20-µL pipette after a brief spin-down, and the pellet was air-dried on ice for 5–10 minutes before being resuspended in nuclease-free water.

#### mRNA Isolation

mRNA was isolated from 1-1.5 mg of total RNA using the NEBNext Poly(A) mRNA Magnetic Isolation Module (New England Biolabs, #E7490L) following the manufacturer’s protocol. After the isolation procedure, the purified mRNA was collected by transferring 88 μL of the supernatant to a 1.5 mL DNA LoBind Tube (Eppendorf, #0030108051). The integrity of the isolated mRNA was assessed using RNA ScreenTape and RNA ScreenTape Sample Buffer (Agilent, #5067-5576 and #5067-5577).

#### RibOxi-seq2 Methods

These were essentially as described^44^ but with a number of modifications.

#### RNA Fragmentation

A 0.25 U/μL benzonase working solution was prepared by mixing 100 μL of 10× benzonase buffer, 899 μL of nuclease-free water, and 1 μL of 250 U/μL Benzonase Nuclease (Millipore Sigma, #E8263). For fragmentation, 88 μL of 10–20 μg mRNA was heated at 95°C for 3 minutes in a ThermoMixer C (Eppendorf) and immediately placed on ice for 3 minutes. Subsequently, 10 μL of 10× benzonase buffer and 2 μL of the benzonase working solution were added to the mRNA solution. The mixture was vortexed, briefly centrifuged, and immediately incubated on ice for 80 minutes. The digestion time requires optimization, particularly for different cell lines and different batches of benzonase.

Post-fragmentation: 100 μL of nuclease-free water and 200 μL of Acid-Phenol:Chloroform (pH 4.5; Invitrogen, #AM9722) were added to the sample. The mixture was vortexed for 15–20 seconds, incubated at room temperature for 2 minutes, and centrifuged at 18,000 × g for 10 minutes at 4°C. The aqueous phase (180–190 μL) was carefully transferred to a new tube. RNA ethanol precipitation was performed as described above. The final RNA was suspended in 64 μL of nuclease-free water. RNA quality was assessed using 1 μL of the sample, confirming the presence of a sharp peak between 25–200 nucleotides, skewing to the right.

#### RNA Oxidation, β-Elimination, and Dephosphorylation

A 200 mM Sodium meta-periodate (NaIO4; Thermo Fisher, #20504) solution was prepared by dissolving 42.78 mg of NaIO4 in 1 mL of nuclease-free water. The solution was protected from light, kept on ice, and used the same day. For the oxidation reaction, 64 μL of fragmented RNA was mixed with 8 μL of oxidation-elimination buffer (pH 8.5) and 8 μL of NaIO4 (200 mM), resulting in a final volume of 80 μL. The mixture was vortexed, briefly centrifuged, and incubated at 37°C with shaking (350 rpm) for 45 minutes in a ThermoMixer C (Eppendorf). Following oxidation, the RNA was purified using ethanol precipitation, and cleaned up using Micro Bio-Spin P-6 Gel Column (Bio-Rad, #7326221), with the RNA eluted in 84 μL of nuclease-free water.

The oxidation-elimination buffer was prepared from 2M L-Lysine monohydrochloride (Sigma-Aldrich, #L5626) in nuclease-free water. The pH was adjusted to 8.5 using 2 M NaOH, and the solution was filtered through a 0.22 μm filter.

For the dephosphorylation reaction, 84 μL of oxidized RNA was mixed with 10 μL of rCutSmart buffer (10×, New England Biolabs, #B6004S), 2 μL of RNaseOUT Recombinant Ribonuclease Inhibitor (Invitrogen, 10777019), and 4 μL of Quick CIP (5 U/μL, New England Biolabs, #M0525S), resulting in a final volume of 100 μL. The reaction was incubated at 37°C for 10 minutes, followed by heat inactivation at 80°C for 2 minutes. The RNA was subsequently purified using ethanol precipitation.

The entire oxidation, β-elimination, and dephosphorylation process was repeated three additional times. Subsequently, RNA cleanup was performed using the Zymo RNA Clean & Concentrator-25 (Zymo Research, #R1017). The final RNA was eluted in 66 μL of nuclease-free water.

Further dephosphorylation reactions involved two stages. In stage I, 66 μL of oxidized RNA was mixed with 8 μL of T4 PNK buffer (10×, pH 6.0), 4 μL of T4 polynucleotide kinase (10 U/μL, New England Biolabs, #M0201), and 2 μL of RNaseOUT inhibitor, resulting in a total volume of 80 μL. The mixture was incubated at 37°C for 3 hours.

In stage II, 80 μL of oxidized RNA from the first reaction was combined with 10 μL of T4 PNK buffer (10×, pH 7.6), 4 μL of T4 Polynucleotide Kinase (10 U/μL, NEB, #M0201S), 20 μL of 10 mM Adenosine 5’-Triphosphate (ATP; New England Biolabs, #P0756S), and 66 μL of nuclease-free water to a final volume of 180 μL. This reaction was incubated at 37°C for 1 hour and inactivated with the addition of 2 μL of 0.5 M EDTA. The RNA was purified by ethanol precipitation and resuspended in 70 μL of oxidation-only buffer. The Oxidation-only buffer was prepared with 4.375mM Sodium Tetraborate Decahydrate (Fisher Scientific, #BP175-500), 50mM Boric Acid (Fisher Scientific, #A74-1), and nuclease-free water. The pH was adjusted to 8.6 using 2 M NaOH, and the solution was filtered through a 0.22 µm filter before use.

An additional oxidation reaction was performed by combining 70 μL of oxidized RNA with 10 μL of NaIO4 solution (200 mM) to a final volume of 80 μL. The mixture was incubated at 37°C with shaking (350 rpm) for 45 minutes. The oxidized RNA was purified using ethanol precipitation. Final cleanup steps included the use of Micro Bio-Spin P-6 Gel Columns and the Zymo RNA Clean & Concentrator-5 (Zymo Research, #R1013), with the RNA eluted in 20 μL of nuclease-free water.

#### 3’-DNA Linker Ligation

The 3’-DNA linker ligation was performed using 20 μL of oxidized RNA, 2 μL of a 3’-DNA linker (10 μM), 2 μL of RNaseOUT inhibitor, 17 μL of 50% PEG 8000, 5 μL of RNA ligase buffer (10×, New England Biolabs, #B0216S), and 4 μL of T4 RNA Ligase 2 truncated KQ (New England Biolabs, #M0373), resulting in a total reaction volume of 50 μL. The reaction mixture was thoroughly mixed, briefly centrifuged, and incubated overnight at 16°C for 18 hours in a thermal cycler. The RNA was subsequently purified using ethanol precipitation and resuspended in 15 μL of nuclease-free water.

#### TBE-Urea Gel Preparation

Gel purification was performed to separate the ligation products from free 3’ DNA linkers.

Two stock solutions were prepared for the TBE-Urea gel. Solution A was made by dissolving 500 g of Urea (Bio-Rad, #1610745) in 500 mL of 40% Acrylamide Solution (Bio-Rad, #1610140) and 100 mL of 10× TBE buffer (Bio-Rad, #1610770). The volume was brought up to 1,000 mL using 1× TBE buffer. Solution B was prepared similarly by dissolving 500 g of Urea in 100 mL of 10× TBE buffer and 500 mL of nuclease-free water, with the final volume adjusted to 1,000 mL using 1× TBE buffer. Both solutions were filtered through a 0.22 μm filter.

A 15% TBE-Urea gel was prepared by mixing 37.5 mL of Solution A, 12.5 mL of Solution B, 400 μL of 10% ammonium persulfate (APS; Bio-Rad, #2610700), and 37.5 μL of TEMED (Bio-Rad, #161-0800) to a final volume of 50.4 mL. The mixture was immediately poured between the glass plates of the Bio-Rad Mini-Protean gel casting apparatus, and a comb was inserted. The gel was allowed to polymerize at room temperature for 60 minutes.

Prior to electrophoresis, 2× RNA loading dye (New England Biolabs, #B0363S) was added to the RNA sample, which was then incubated at 72°C for 2 minutes and placed on ice for 3 minutes. The samples were loaded, and electrophoresis was performed at 20W for 50 minutes.

The gel was stained with SYBR Gold Nucleic Acid Gel Stain (Invitrogen, #S11494) by incubating it in a 1× TBE staining solution for 15 minutes with gentle shaking. RNA bands were visualized using a Dark Reader Non-UV Transilluminator (Model: DR-46B), and the desired fragment was excised and transferred to a tube. Images of the gel were captured both before and after excision.

#### Acrylamide Gel RNA Extraction

To extract RNA product from the acrylamide gel, LoBind tubes were prepared with 1.5 μL GlycoBlue Coprecipitant and 33 μL Sodium Acetate.

A hole was pierced in the bottom of a 0.6 mL tube using an 18G needle, holding the needle near the tip and carefully pressing it into the tube’s bottom. The 0.6 mL tube was placed inside a 2 mL tube, and the gel slices were added to the 0.6 mL tube. The assembly was centrifuged at maximum speed for 1 minute at 4°C, allowing the crushed gel to flow into the larger tube.

Each tube was supplemented with 300 μL of nuclease-free water, and the mixture was incubated in a ThermoMixer C (Eppendorf) at 70°C for 10 minutes with shaking at 1800 rpm. This incubation was repeated an additional two times. Following incubation, the tubes were briefly centrifuged, and the supernatant was transferred to a Costar SpinX Centrifuge Tube Filter (0.45 μm; #8162). The SpinX tube was centrifuged at 10,000 rpm for 1 minute, and the flow-through was transferred into prepared LoBind tubes containing GlycoBlue and sodium acetate. Ethanol precipitation was performed, and the RNA product was eluted in 22 μL of nuclease-free water.

#### 5’-RNA Linker Ligation

To perform the 5’-RNA linker ligation, 2 µL of 5’-RNA linker (50 µM) was denatured at 72°C for 2 minutes and immediately placed on ice. A ligation reaction mixture was prepared by combining 22 µL of the ligated RNA product, 4 µL of 100% DMSO, 4 µL of T4 RNA Ligase Reaction Buffer (10×, B0216S), 4 µL of ATP (10 mM), and 1 µL of RNaseOUT inhibitor, resulting in a total volume of 35 µL. Next, 35 µL of the prepared ligation mixture was combined with 3 µL of T4 RNA Ligase 1 (New England Biolabs, #M0204S) and 2 µL of the 5’-RNA linker (50 µM), bringing the final reaction volume to 40 µL. The reaction was mixed, briefly centrifuged, and incubated at 25°C for 1 hour. Following the incubation, RNA purification was carried out using the Monarch Spin RNA Cleanup kit (New England Biolabs, #T2030) according to the manufacturer’s instructions, and the RNA product was eluted in 20 µL of nuclease-free water.

#### cDNA Synthesis

For cDNA synthesis, 20 µL of the ligated RNA product was combined with 1 µL of 10 µM Reverse Transcriptase (RT) primer, 2 µL of 10 mM dNTPs, and 2 µL of nuclease-free water in a final volume of 25 µL. The mixture was denatured at 65°C for 5 minutes, immediately placed on ice, and briefly centrifuged. To this, 8 µL of ProtoScript II buffer (5×, New England Biolabs, #B0368S), 1 µL of RNaseOUT inhibitor, 4 µL of 0.1 M DTT (New England Biolabs, #B1034A), and 2 µL of ProtoScript II Reverse Transcriptase (New England Biolabs, #M0368S) were added to bring the final reaction volume to 40 µL. The reaction was incubated at 50°C for 1 hour to synthesize the first strand of cDNA. To hydrolyze the RNA, 4 µL of 1 N Sodium Hydroxide Solution (Fisher Chemical, #SS266-1) was added, and the mixture was incubated at 95°C for 15 minutes. The reaction was neutralized with 4 µL of 1 M Tris–HCl (pH 7.5; Invitrogen, #15567027), and the cDNA was purified using 1.8× AMPure XP Reagent (Beckman Coulter, #A63881) following the manufacturer’s protocol.

#### cDNA Purification Using AMPure XP Reagent

The cDNA was purified by adding 1.8× AMPure XP reagent to the cDNA sample (48 µL cDNA + 86.4 µL beads). The mixture was thoroughly mixed by pipetting up and down 10 times and incubated at room temperature for 10 minutes. The tubes were briefly spun down and placed on a magnetic stand for 5 minutes until the liquid became clear. The supernatant was carefully removed and discarded, and the DNA-bound beads were washed twice with 500 µL of freshly prepared 80% ethanol. After each wash, the beads were incubated on the magnetic stand for 30 seconds, and the supernatant was discarded. Residual ethanol was carefully removed with a pipette, and the beads were air-dried for 5–10 minutes. The dried beads were resuspended in 32 µL of nuclease-free water, incubated at room temperature for 2 minutes, and briefly spun down. The eluate (30 µL), containing the purified cDNA, was carefully transferred to a new tube for subsequent use.

#### Library Amplification and Library Quality Control (QC)

Library amplification was performed using dual-index sequences. The reaction was set up with 30 µL of cDNA, 10 µL of 5× Q5 Reaction Buffer, 1 µL of 10 mM dNTPs, 2.5 µL of 10 µM i7 index, 2.5 µL of 10 µM i5 index, 0.5 µL of Q5 High-Fidelity DNA Polymerase (New England Biolabs, #M0491), and 3.5 µL of nuclease-free water, in a final volume of 50 µL. Thermocycling conditions were as follows: initial denaturation at 98°C for 30 seconds, followed by three cycles of 98°C for 10 seconds, 52°C for 30 seconds, and 72°C for 30 seconds. This was followed by 18 cycles of 98°C for 10 seconds, 62°C for 30 seconds, and 72°C for 30 seconds. The final extension was performed at 72°C for 2 minutes, and the reaction was held at 4°C. The amplified library was purified using ethanol precipitation and P-6 columns, followed by size selection using SPRIselect beads (Beckman Coulter, #B23318) to ensure high-quality libraries.

#### SPRI-Based Size Selection

SPRI-based size selection was performed using SPRIselect (Beckman Coulter, #B23318) to ensure appropriate library fragment size distribution. For left-side selection, 60 µL of SPRIselect beads (1.2× ratio) was added to a 50 µL amplified library sample. The beads were thoroughly mixed by pipetting up and down 10 times and incubated at room temperature for 1 minute. The reaction tube was then placed on a magnetic stand for 5–10 minutes to allow the beads to settle. Once the liquid clarified, the supernatant was carefully discarded. The beads were washed twice with 500 µL of freshly prepared 85% ethanol. After each wash, the supernatant was removed following a 30-second incubation, and the beads were air-dried for 5–10 minutes to ensure complete ethanol removal. The purified DNA was eluted by resuspending the beads in 52 µL of nuclease-free water. After incubation for 1 minute and magnetic separation, the eluate (50 µL) was carefully collected, leaving the beads behind.

For right-side size selection (0.7×/1.1×), 35 µL of SPRIselect beads (0.7× ratio) was added to the 50 µL library sample and mixed thoroughly. Following incubation and magnetic separation, the supernatant containing the desired DNA fragment was transferred to a new reaction tube, and the discarded beads contained larger fragments. To the supernatant, 55 µL of SPRIselect beads (1.1× ratio) was added to remove smaller fragments. After incubation, magnetic separation, and two ethanol washes, the beads were air-dried, and the DNA was eluted in 22 µL of Resuspension Buffer (Illumina, #15026770). The final eluate containing the size-selected library was carefully transferred to a new LoBind tube and stored at −80°C for subsequent analysis.

#### RibOxi-seq2 Library Preparation and Sequencing

Library size distribution was confirmed using the D1000 ScreenTape and reagents on the 4200 TapeStation system (Agilent, #5067-5582 and #5067-5583). Library concentration was quantified using the Qubit dsDNA BR Assay Kit (Invitrogen, #Q32853) on a Qubit 2.0 fluorometer (Invitrogen). For sequencing, libraries were initially run on an Illumina MiSeq system using the MiSeq Reagent Nano Kit v2 (300 cycles; Illumina, #MS-103-1001) with a loading concentration of 9 pM. For higher throughput sequencing, libraries were diluted to a final concentration of 1050 pM and sequenced on an Illumina NextSeq 2000 system using the NextSeq 2000 P2 Kit (200 cycles; Illumina, #20046812). A sequencing depth of 30 million reads per sample was achieved to ensure sufficient coverage for downstream analysis.

#### RibOxi-seq2 analysis

RibOxi-seq2 data were processed following the previously established pipeline (https://github.com/yz201906/RibOxi-seq, https://github.com/xxy103/RibOxi-seq) ^44^. In brief, sequencing adapters were trimmed using Cutadapt v2.7 (https://github.com/marcelm/cutadapt). Paired-end reads were subsequently merged into single-end reads with PEAR v0.9.11. Unique molecular identifiers (UMIs), consisting of 10 randomized nucleotides, were extracted and appended to the fastq headers using the “move_umi.py” script. Quality-filtered reads (with length longer than 20nt, Q-value over 20) were then mapped to the reference genome GRCh38_no_alt_analysis_set_GCA_000001405.15.fasta using STAR v2.7.1a. The aligned reads were filtered and de-duplicated using UMI-TOOLS v1.1.1. Single-nucleotide 3’-end coverage profiles indicative of Nm sites were calculated using Samtools v1.9 and Bedtools v2.29.0.

#### rRNA, snRNA, and tRNA

rRNA and snRNA sequences were retrieved from https://rnacentral.org/, tRNA sequences were downloaded from https://gtrnadb.org/.

#### Snoscan RNA modification prediction

The annotated snoRNA sequences were downloaded from Human snoDB ^30^, the top190 expressed human snoRNAs in 5 human cell lines (HepG2, HEK293T, PC3, A549, MDA-MB-231) were obtained from previous publication ^68^. Putative RNA modification sites within these snoRNAs were predicted with snoScan v1.0 ^67^, using as reference the Nm regions (± 7nt) identified through RibOxi-seq2 analysis.

#### SnoRNA-RNA interactions

The snoRNA-RNA base pairing was computed with snoDB online tools (https://bioinfo-scottgroup.med.usherbrooke.ca/snoDB/sequence_similarity_search/)^30^, predicted interactions were filtered based on alignment scores and thermodynamic plausibility. The RNA-RNA interactions Minimum Folding Energy (MFE) was determined using RNAduplex (ViennaRNA v2.5.1) ^104^, which computes the MFE of hybridization between snoRNA and Nm regions (± 7nt) identified from RibOxi-seq2 analysis.

#### Metagene analysis

Metagene plot summarizing the mRNA features were generated with metaPlotR. (https://github.com/olarerin/metaPlotR)

#### RiboMeth-Seq

H9 hESCs were maintained on Matrigel-coated (Corning, #354277) plates with Essential 8 media (Thermo Fisher, #A1517001). Stem cells were differentiated to neurons for 14 days per the i^3^ neuron protocol^105^. hESCs were prepared for differentiation once 70-80% confluent. Differentiated cells were manually removed. Stem cells were released from the plate with Accutase (Sigma-Aldrich, #SCR005) incubation. Media was added to the cell solution and cells were centrifuged. Cells were resuspended in induction media (IM) supplemented with 2µg/ml doxycycline (Dox) (Milipore Sigma, #D9891) and 10µM Rock Inhibitor (Tocris, #1254). IM was made with DMEM/F12 with HEPES (Thermo Fisher, #11330032), N2 supplement (Thermo Fisher, #17502048), MEM Non-essential amino acids (Thermo Fisher, #11140050), and L-Glutamine (Thermo Fisher, #25030081). Cells were plated at 1.5×10^5^ cells/well in a 6-well plate. Cells were fed daily with IM and received 2 additional days of Dox treatment. Cells were then replated in IM on poly-D-lysine/laminin-coated 6-well plates (Millipore Sigma P1149, Fisher Scientific, #23-017-015) at 5×10^5^ cells/well. The day after plating, the media was fully changed to cortical neuron culture media (CM) which is composed of DMEM/F12 with HEPES and Neurobasal medium (Thermo Fisher, #21103049) supplemented with laminin, BDNF (Thermo Fisher, #450-02-10UG), GDNF (Thermo Fisher, #450-10-10UG), NT3 (Thermo Fisher, #450-03-10) and B27 supplement (Thermo Fisher, #17504044). Subsequently, half media changes with CM were performed on every other day until the 14^th^ day of differentiation. Cells were rinsed with PBS and then were collected in TRIzol Reagent (Thermo Fisher, # 15596018). 0.2ml of chloroform per 1ml of TRIzol was added and samples were centrifuged. The aqueous phase was transferred to a new tube and 0.5ml of isopropanol was added. Samples were incubated at −80°C for 10 minutes and then centrifuged to pellet the RNA. The pellet was washed with 75% ethanol and resuspended in nuclease-free water.

Samples were prepared in triplicate for RiboMeth-Seq following ^42^ with some modifications. Briefly, 250ng RNA per sample was subjected to alkaline hydrolysis. Fragmented RNA was then ethanol precipitated and run on a TapeStation to check size distribution. Samples were 3’ end dephosphorylated by incubating for 3 hrs at 37°C with T4 PNK (New England Biolabs, M0201S) and pH6 T4 PNK buffer. Then 5’ end phosphorylation was achieved by adding additional T4 PNK, T4 PNK reaction buffer, and ATP and incubating for 30 minutes. A 3’-DNA linker was ligated to the RNA with T4 RNA ligase 2 truncated KQ (NEB, M0373S). Then a 5’-RNA linker was ligated to the RNA. The linker was denatured and then the ligation reaction was prepared utilizing T4 RNA ligase 1 (NEB, M0204S. cDNA was synthesized using Superscript III (Thermo Fisher, #18080044). The RNA was hydrolyzed and the cDNA purified with AmpureXP beads (Beckman Coulter, A63880). Libraries were amplified with KAPA HiFi DNA polymerase (Roche, #07958927001) and sequenced on Illumina NovaSeq. Adaptor sequences, reverse transcription primers, and PCR primers were applied according to the procedures described above.

RiboMeth-Seq Analysis by MethScore was calculated as described^40,106^. The 2 nucleotides flanking each position were used as the flanking region. Briefly, across all positions, the sum of 3’-ends’of profile at the n position and the 5’-ends of profile at the n+1 position was calculated. For each position, the sum of ends was divided by ½ of the sum of the weighted end sum for the flanking positions divided by the sum of the weights. This was subtracted from 1 to give the MethScore. Standard deviation was calculated across 3 replicates.

#### Nm-VAQ Quantitation

500 ng of total RNA from CT2-NGN2 neuron cells was mixed with 50 pmol of the RNA/DNA chimera. The volume was adjusted to 11 µl using 10 µM Tris pH 7.0 buffer. For mRNA-specific targeting, 500 ng of polyA-selected RNA (NEB, #E7490L) was used. Samples were incubated at 95°C for 1 minute and immediately transferred to ice. Subsequently, 5 µl of the annealed RNA/chimera mixture was combined with 1 µl of RNase H enzyme (NEB, #M0297S), 1 µl of 10x RNase H buffer (NEB, #B0297S), and 3 µl of nuclease-free water. The remaining 5 µl of the annealed RNA/chimera mixture was mixed with 1 µl of 10x RNase H buffer and 4 µl of nuclease-free water. The samples were thoroughly mixed by pipetting and incubated at 37°C for 30 minutes.

At this stage, the protocol diverges for highly abundant rRNA and low abundance mRNA. For highly abundant RNAs, the samples were incubated at 90°C for 10 minutes to denature the RNase H enzyme, and then placed on ice. After denaturation, the samples were diluted 1:5. Then, 1 µl of the diluted sample was used for cDNA synthesis with SuperScript III Reverse Transcriptase (Invitrogen, #18080044) and random hexamers. Subsequently, 1 µl of cDNA was used for RT-qPCR with PowerSybr (Applied Biosystems, #4368706).

For low abundance mRNA, the reaction volume after the 30-minute RNase H cleavage step was increased to 30 µl with sterile nuclease-free water, and 30 µl of Phenol-chloroform-isoamyl alcohol mixture (Millipore Sigma, #77617) was added. After vigorous vortexing, the mixture was centrifuged at 12,000 g for 5 minutes. Approximately 20 µl of the upper aqueous phase was transferred to a clean 1.7 mL Eppendorf tube. Cytiva Microspin G-50 columns (Cytiva, #27533001) were prepared by loosening the cap, removing the bottom plug, and placing them into 2 mL collection tubes. Excess buffer was removed by centrifuging at 700 g for 1 minute. The column was then transferred to a clean 1.7 mL Eppendorf tube, and the upper phase from the previous extraction was added to the column. To elute, the tube was centrifuged at 700 g for 2 minutes. 1.5 µl of the eluate was used for cDNA synthesis with SuperScript III and random hexamers. Finally, 1 µl of the cDNA was used for RT-qPCR with PowerSybr.

### QUANTIFICATION AND STATISTICAL ANALYSIS

All statistical tests are described in corresponding figure legends and methods section.

---

**Figure S1**. RibOxi-seq2 peaks in tRNA, snRNA, RN7SK from induced neurons. Modified nucleotides annotated in reference are highlighted. The RN7SK Nm site was quantified at 75% using Nm-VAQ.

**Figure S2**. Some additional representative examples of Nm sites in mRNAs, illustrating similar modification profiles between dirrerent cells. Percentages adjacent to some peaks denote modification levels quantified by Nm-VAQ.

**Figure S3**. Box plot comparison of minimum folding energy (MFE) across snoRNAs and RNA regions encompassing each Nm site (±7 nt). For each snoRNA species, the potential MFE values are shown; boxes denote the 25th–75th percentiles with a line at the median, and whiskers indicate 1.5× the interquartile range.

**Figure S4.** Representative UCSC genome browser images illustrating bigWig tracks from CT2 and H9 induced neurons as well as small-deletion induced neurons and, in almost all cases, 293T cells. Nm positions are denoted by RibOxi-seq2 peaks, while U3 snoRNA CLASH fragment overlaps from HEK293T cells are shown at the top of each panel.

**File S1.** Nm sites in 18S and 28S rRNA identified by RibOxi-Seq2 and RiboMeth-Seq, related to Figure 2.

**File S2**. Nm sites in CT2 and H9 cell lines identified by RibOxi-Seq2, related to Figure 3.

**File S3.** Nm-VAQ quantifications, related to Figures 4, 6, S2 and STAR Methods.

**File S4**. Nm-VAQ oligos and qPCR-primers, related to Figures 4, 6, S2 and STAR Methods.

