## Supplementary figures and images for "Genome-wide profiling of RNA 2’-*O*-methylation in neurons and identification of orphan snoRNA targets"

### Supplemental Figure 1

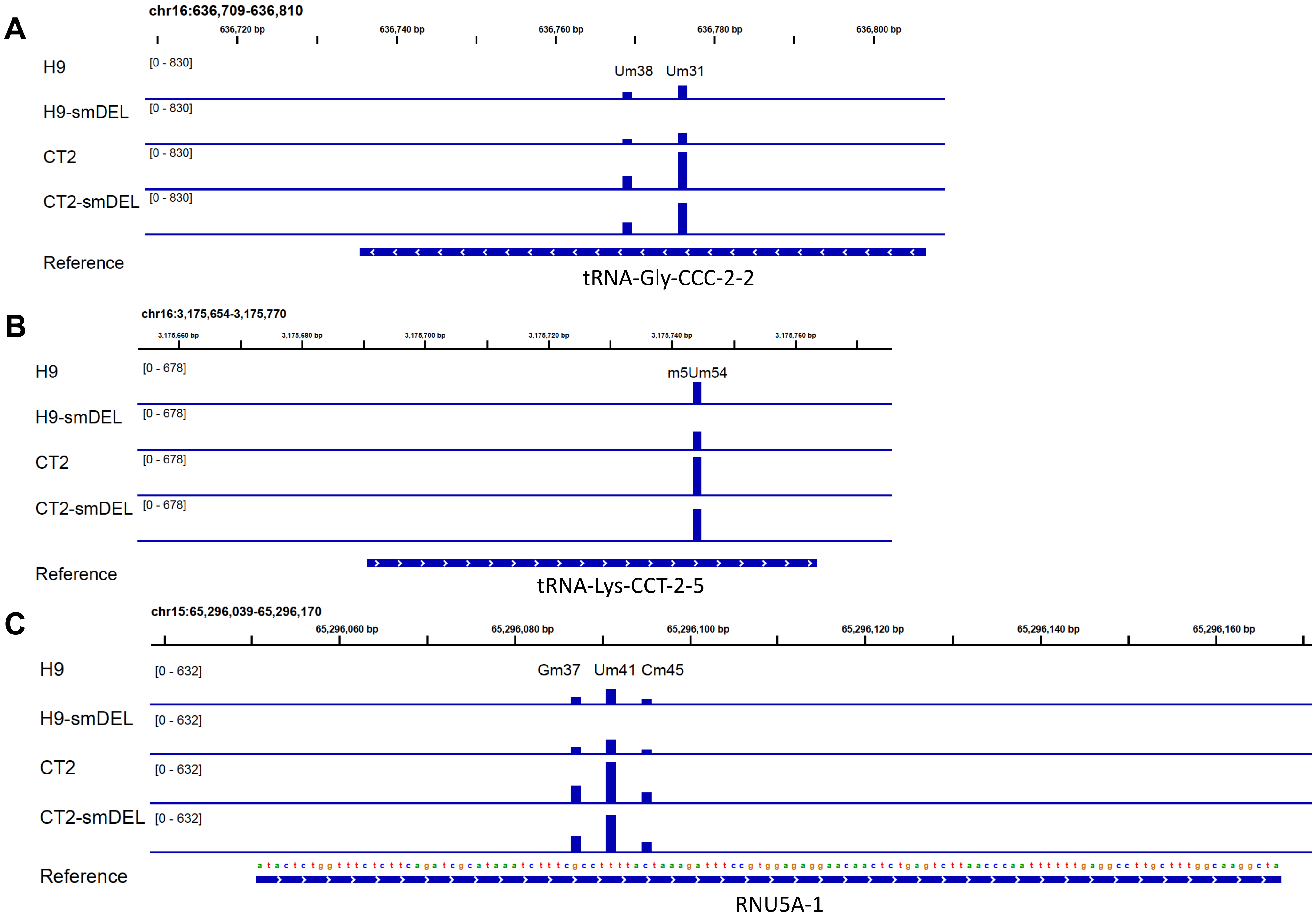

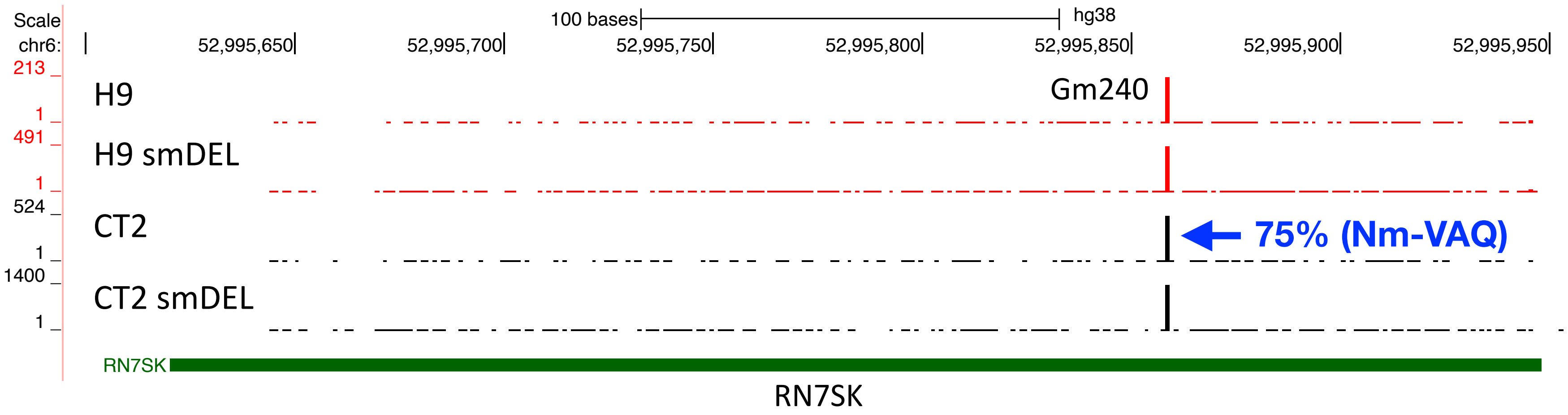

### Supplemental Figure 2

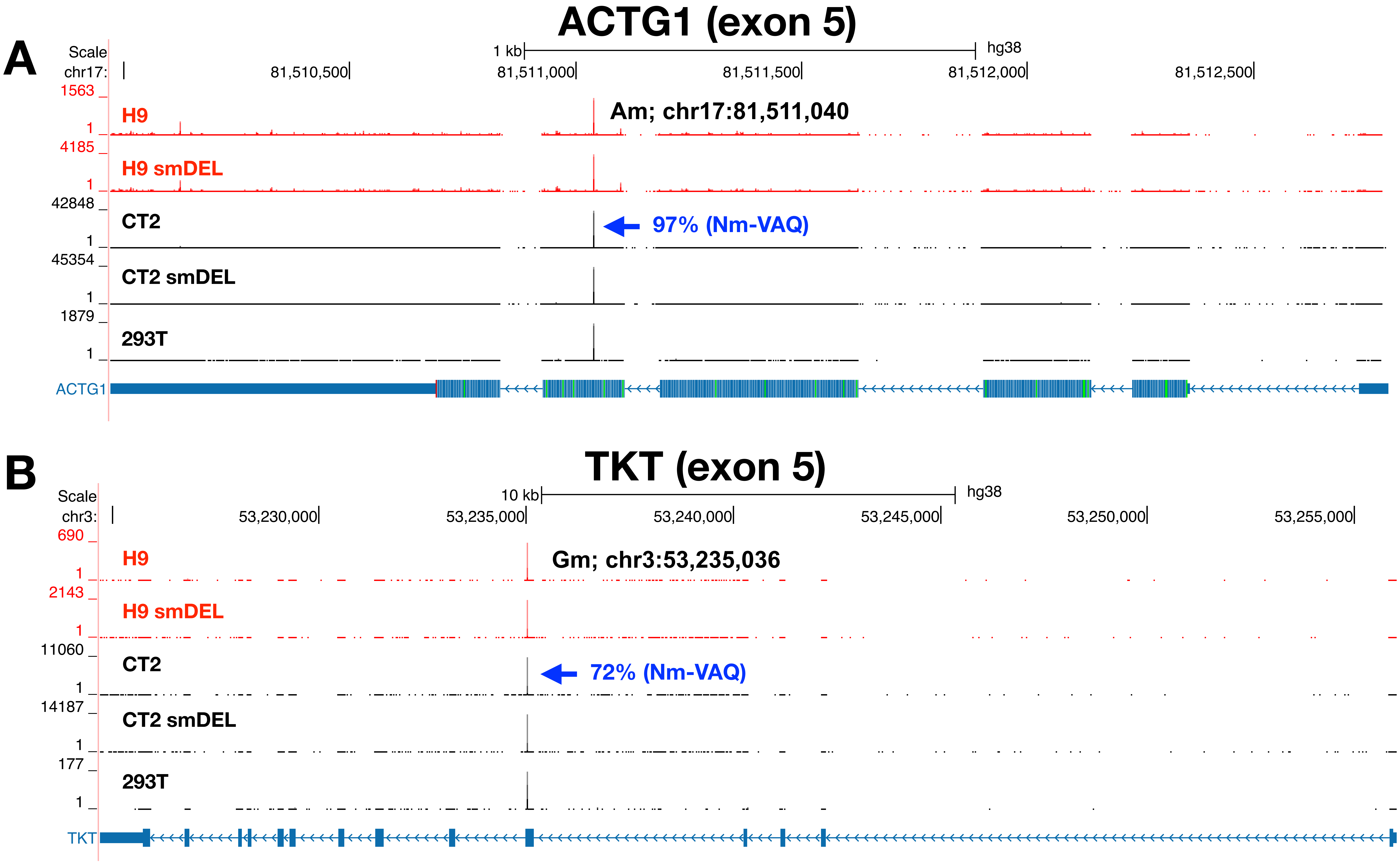

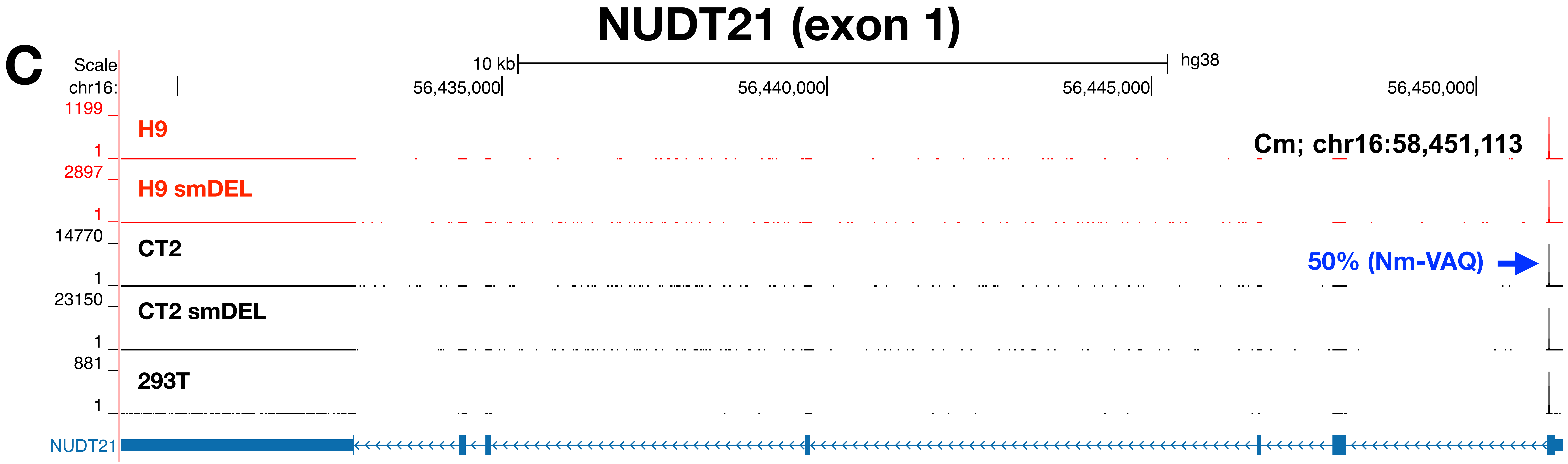

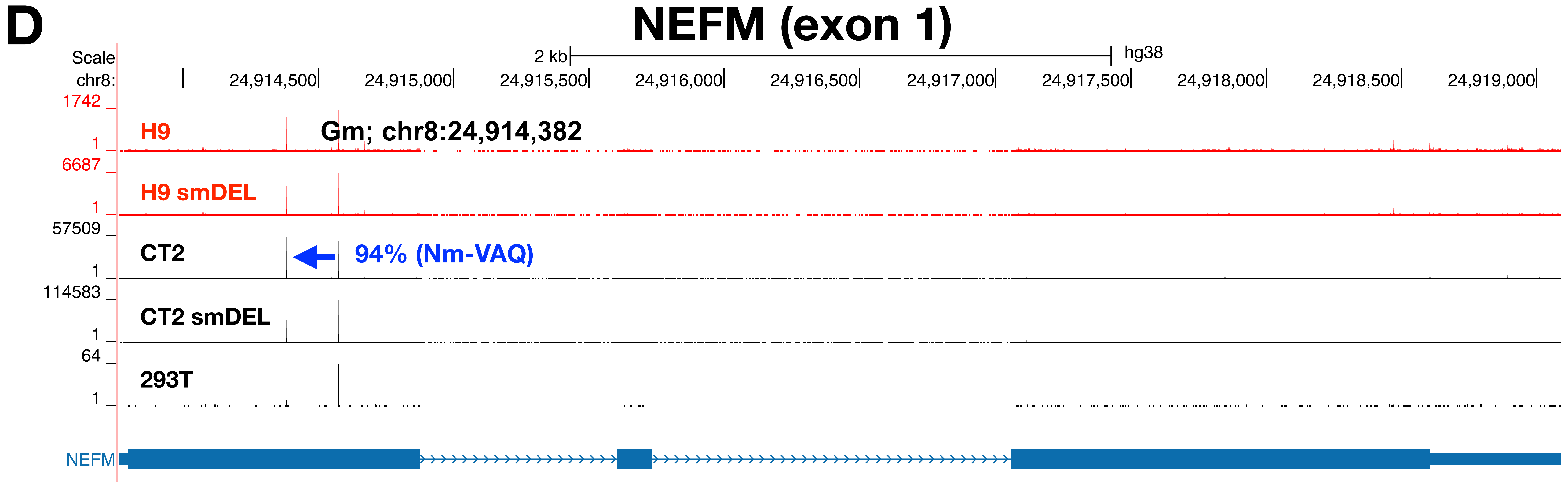

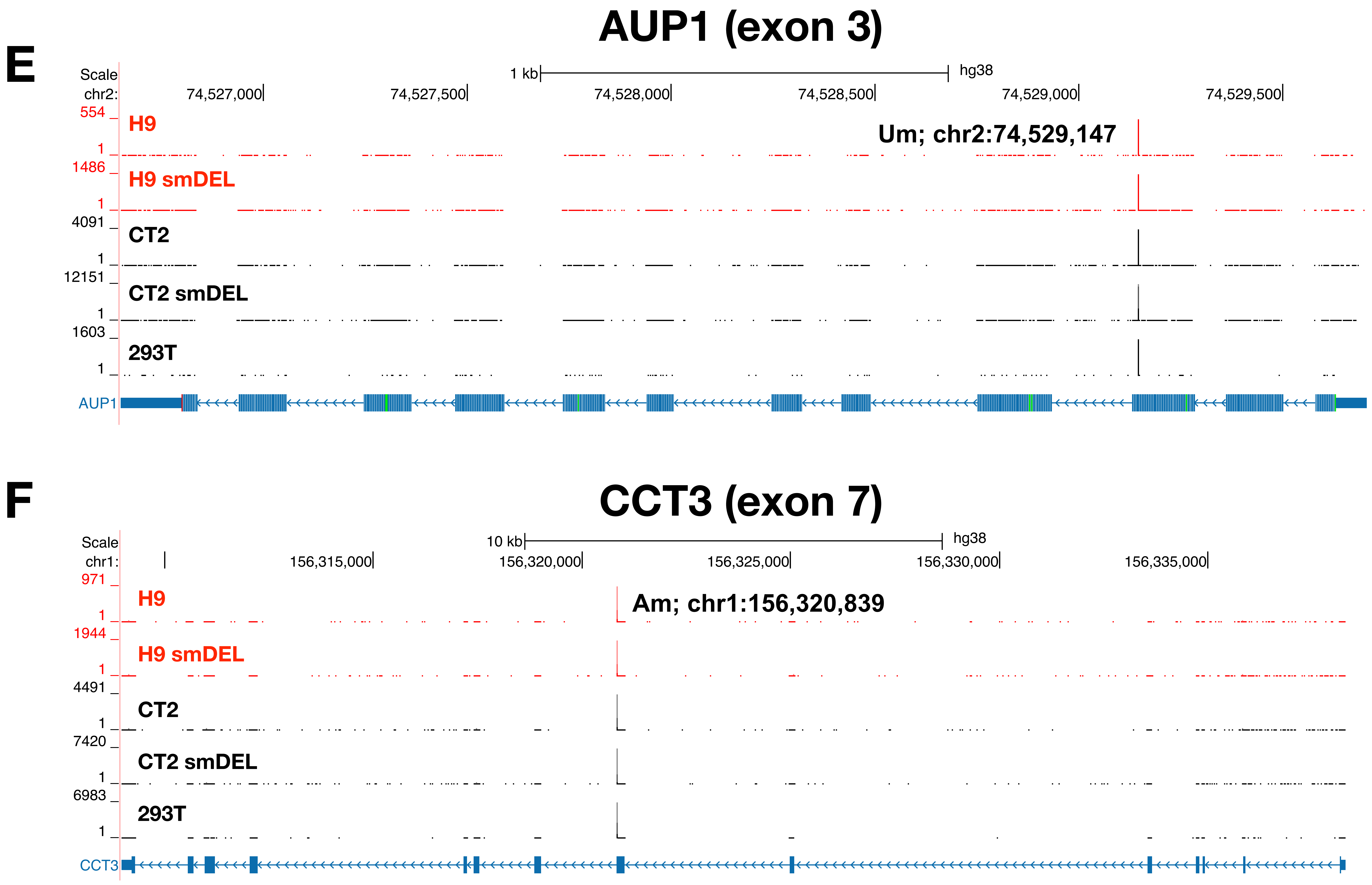

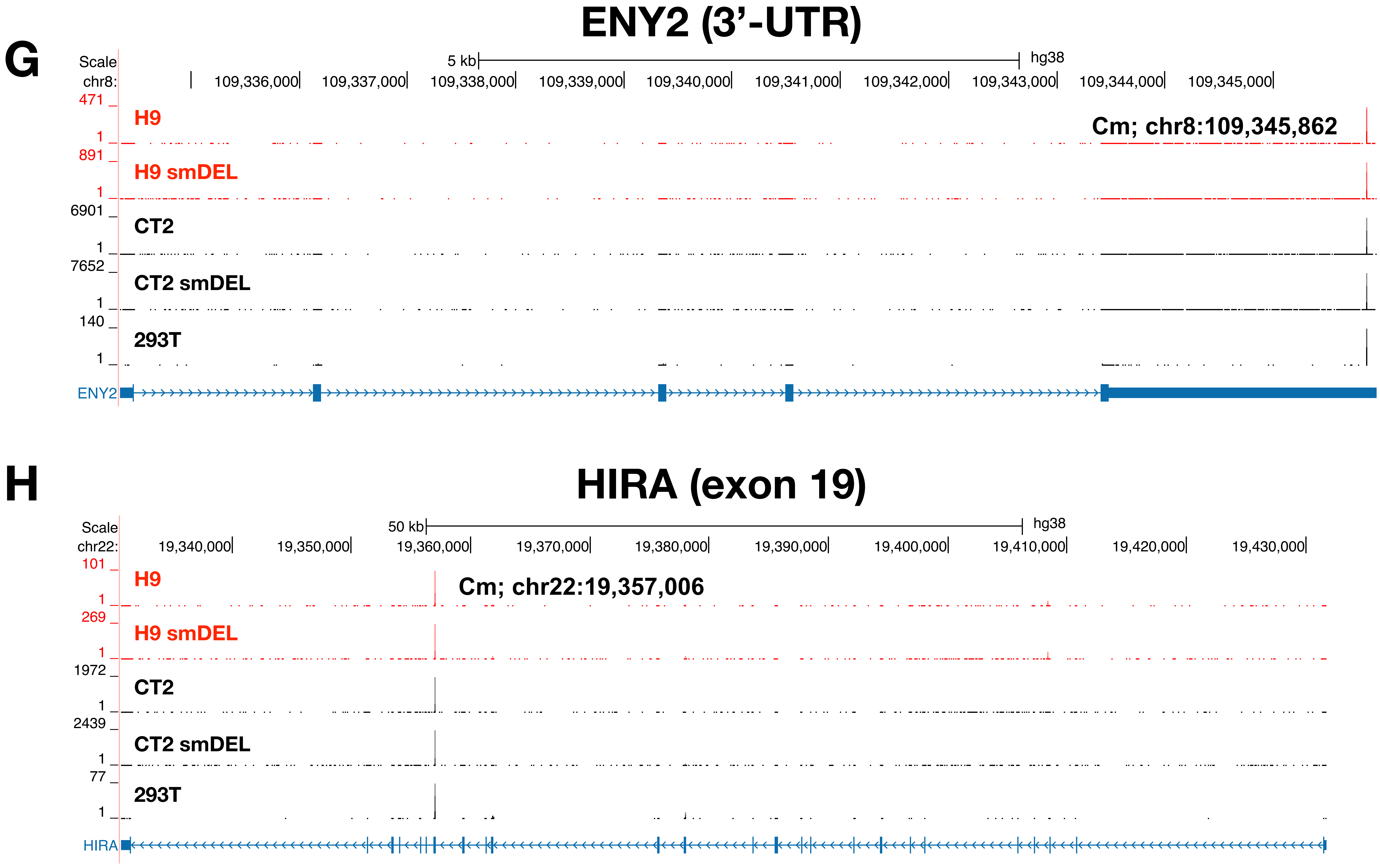

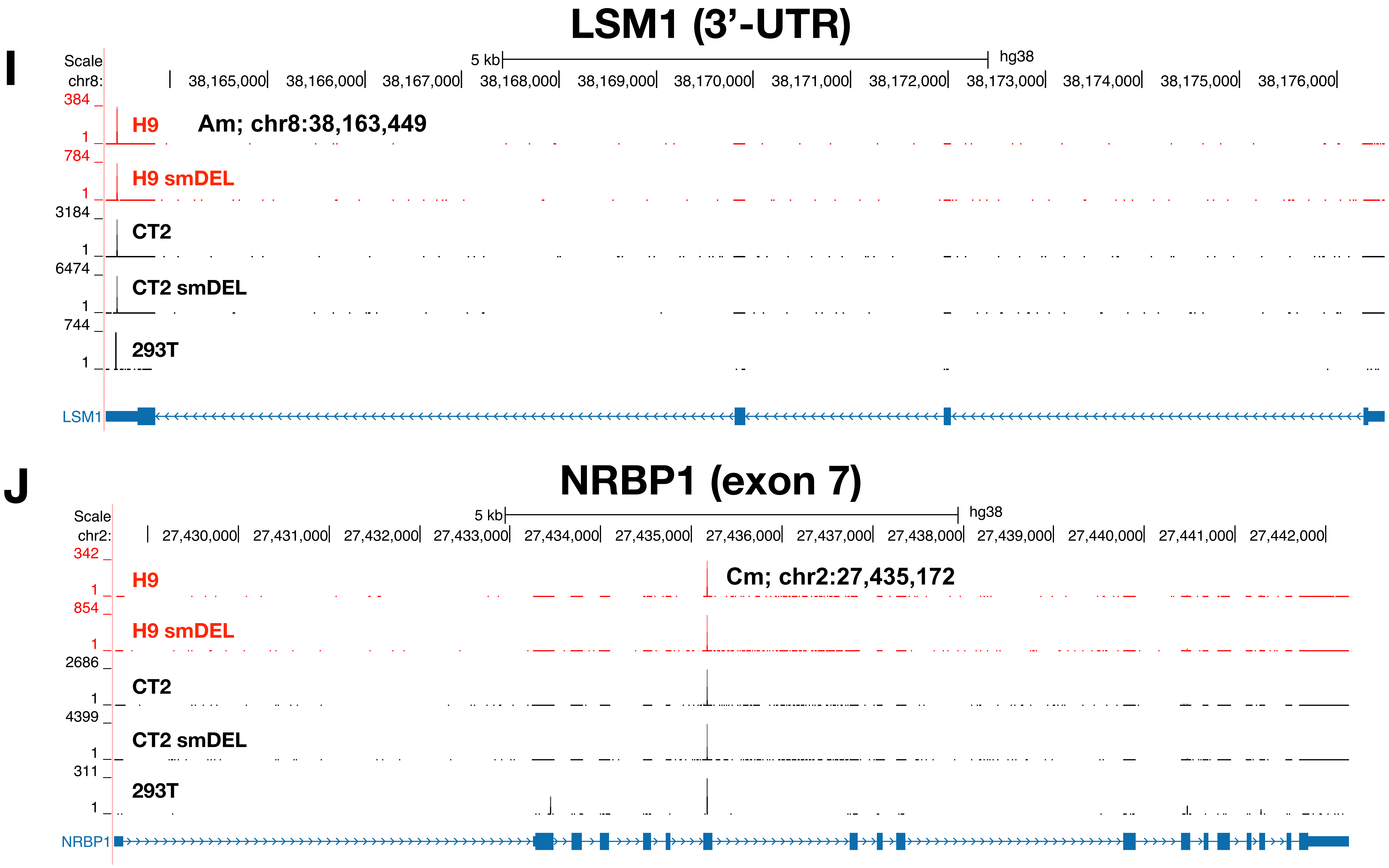

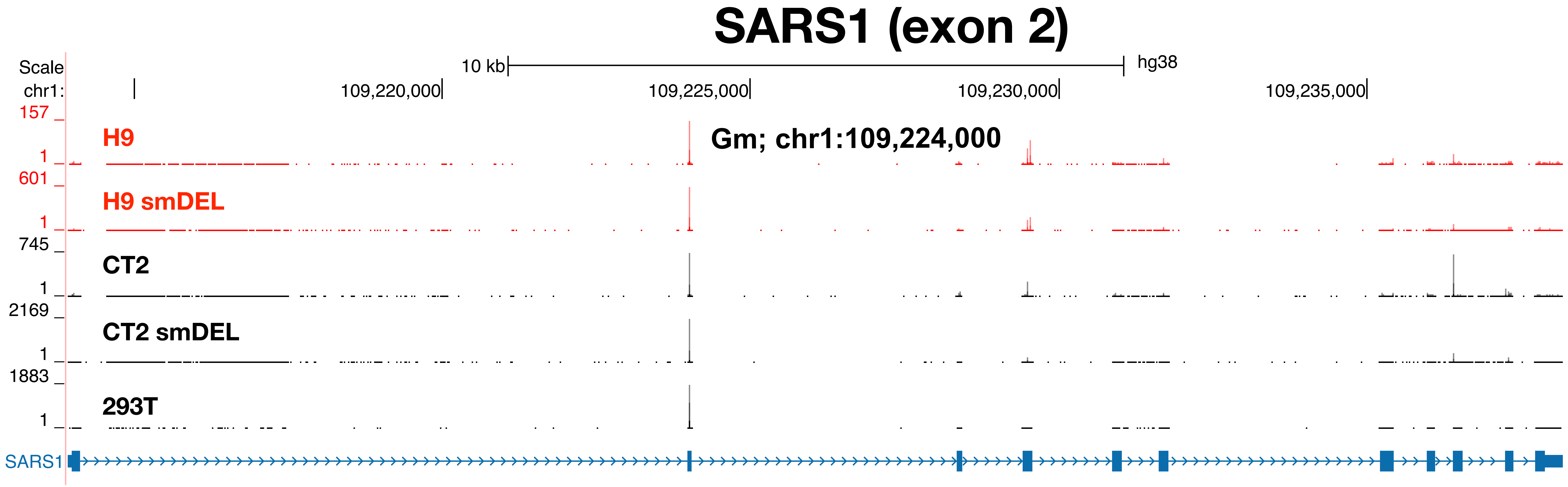

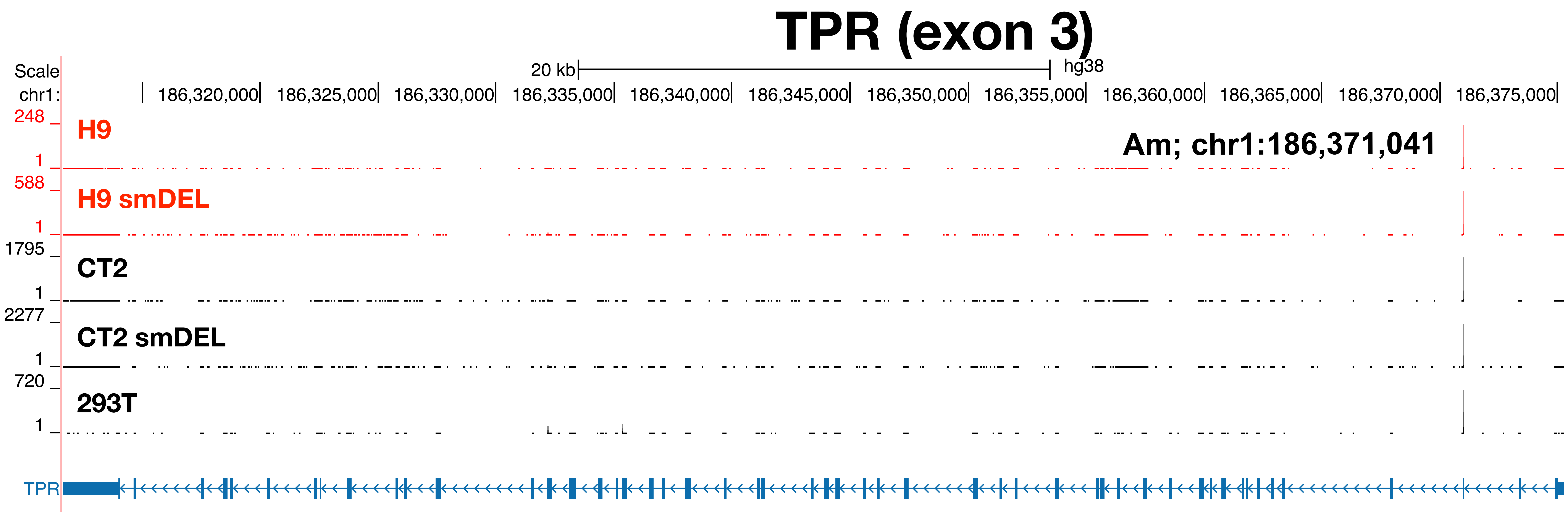

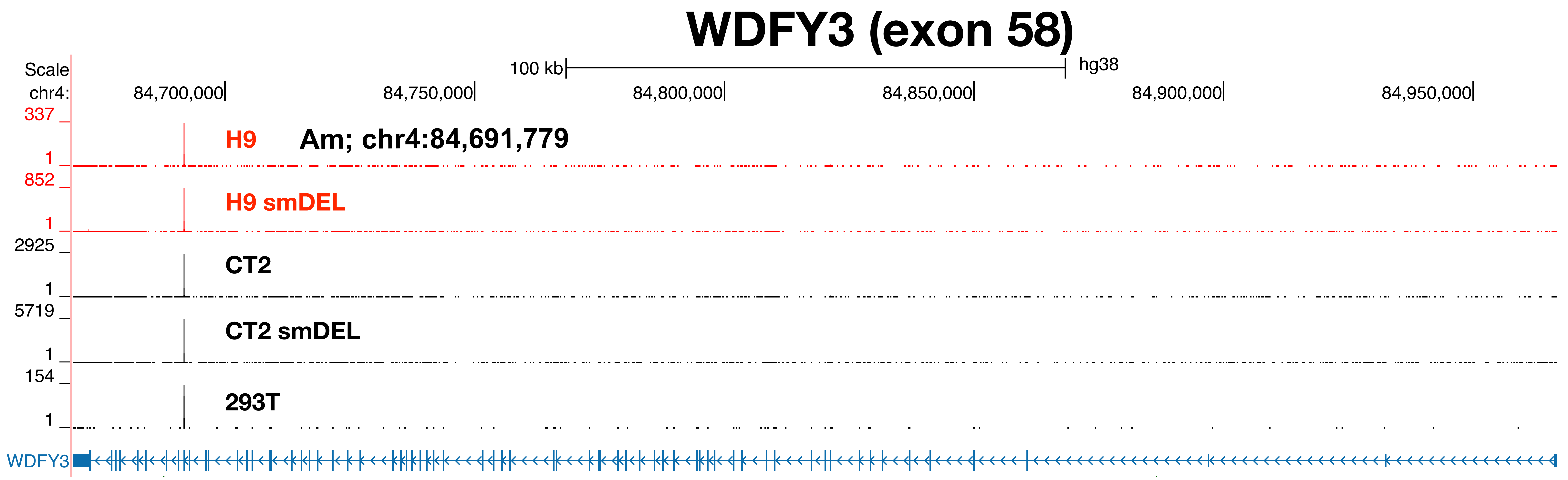

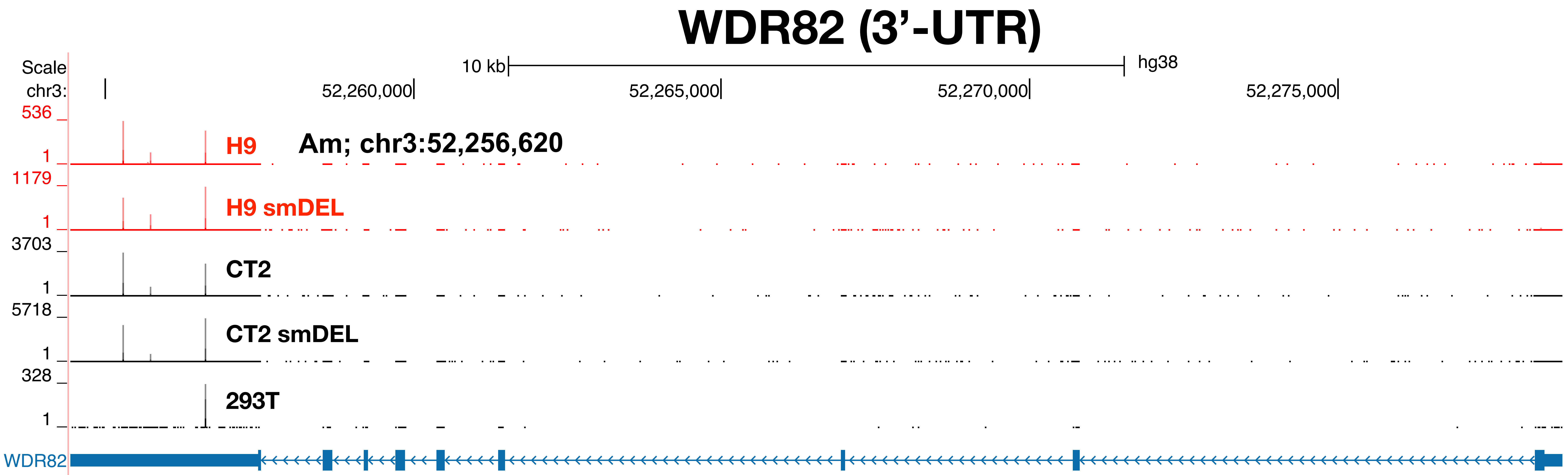

### Supplemental Figure 3

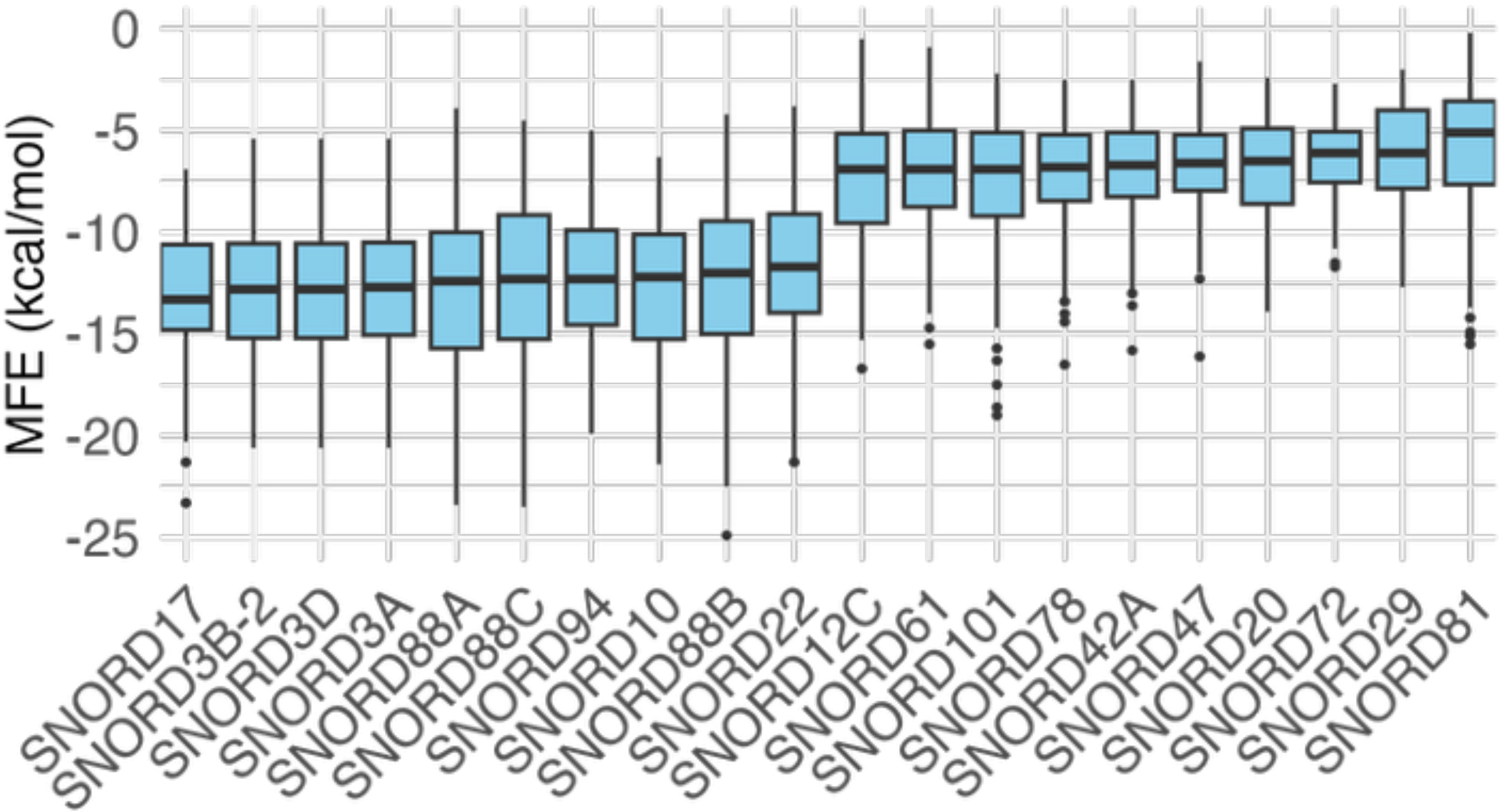

### Supplemental Figure 4

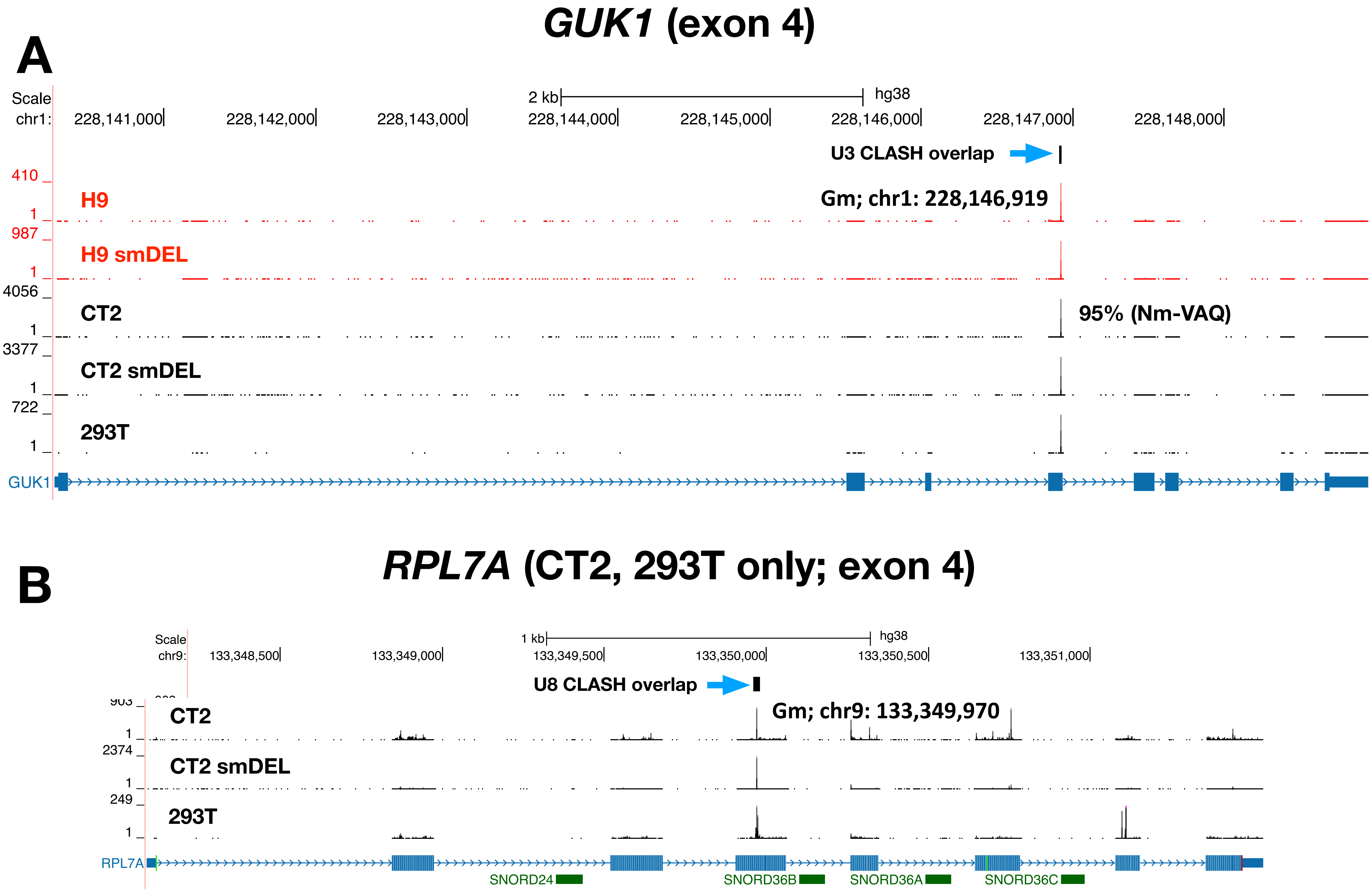

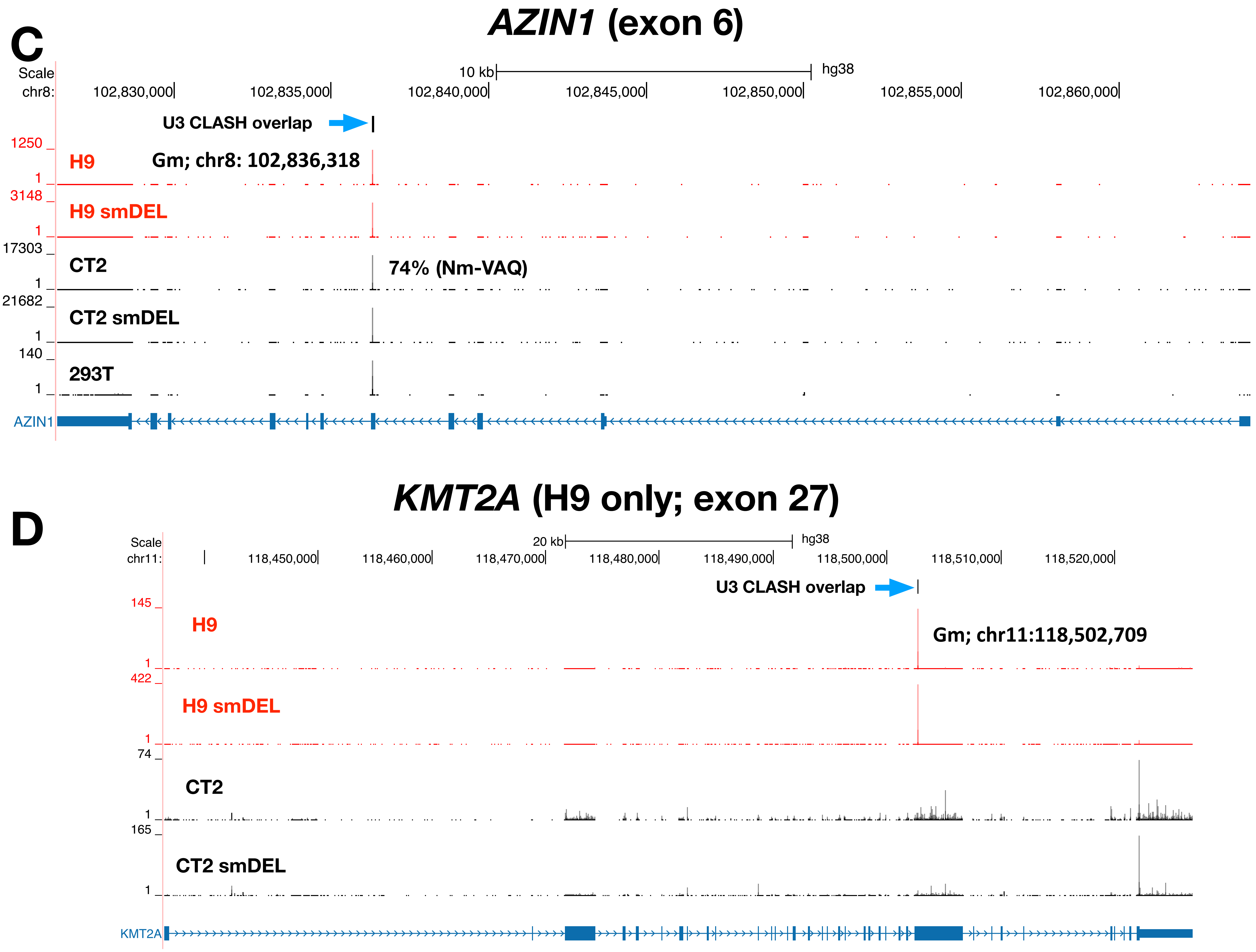

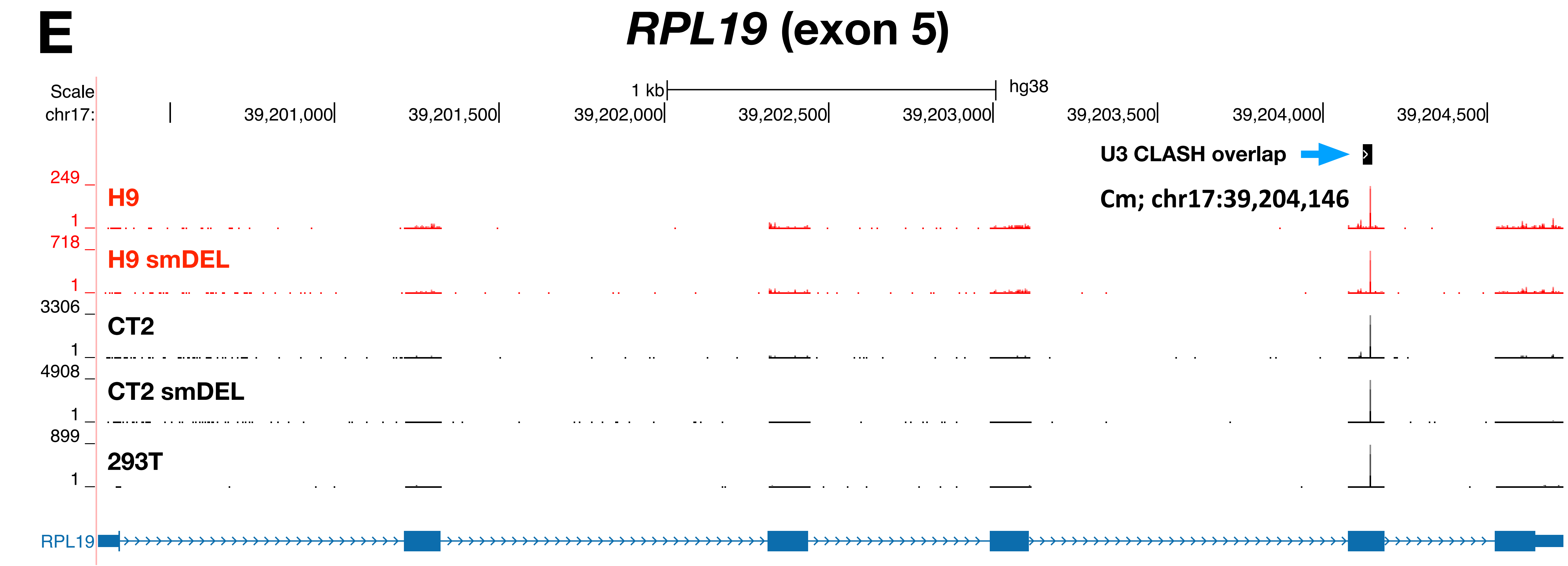

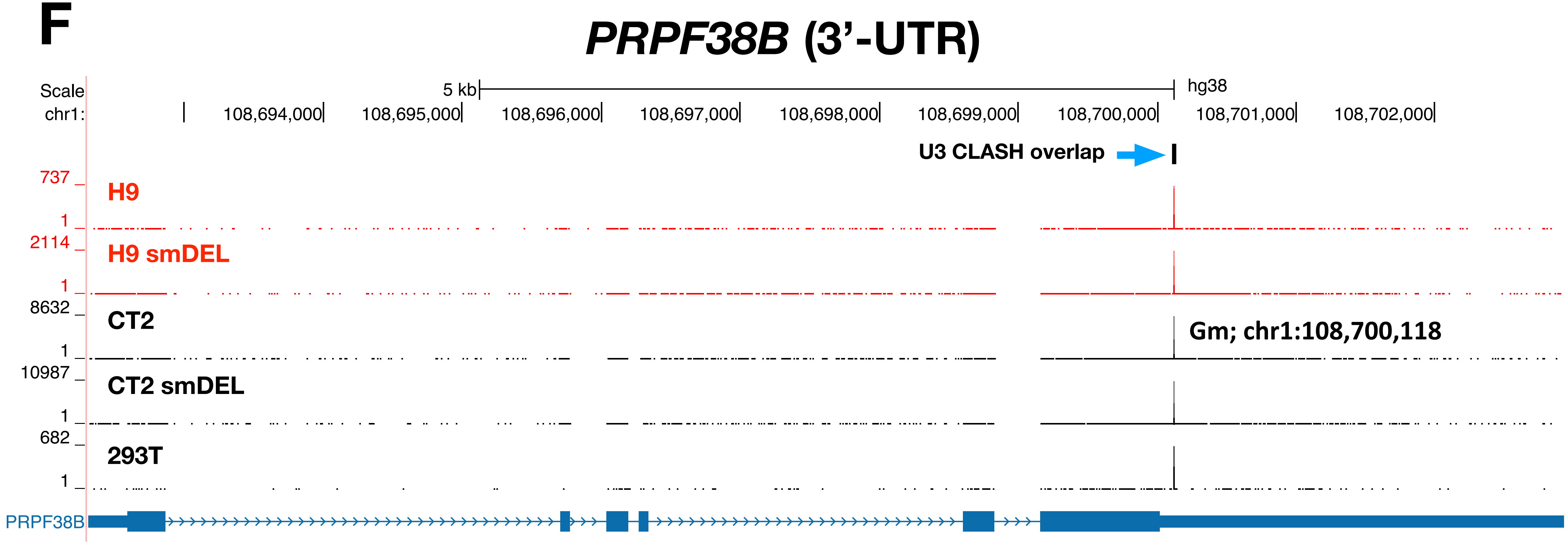

**G**

#### *EIF4G1* (exon 3)

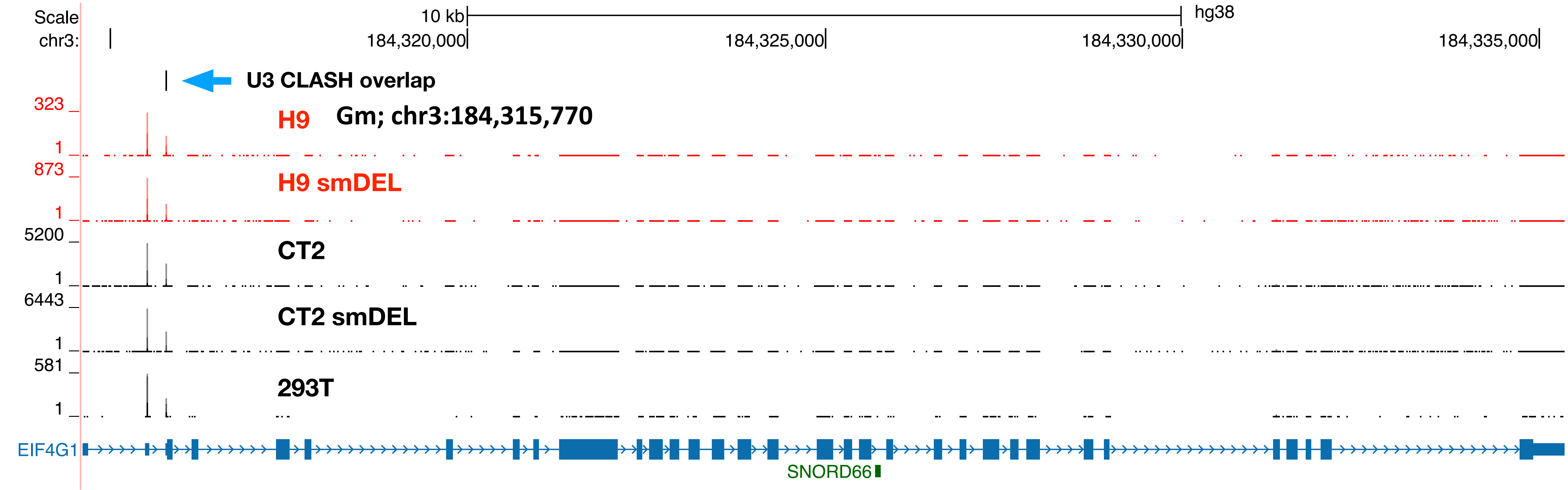


**H**

#### *EIF3F* (exon 7)

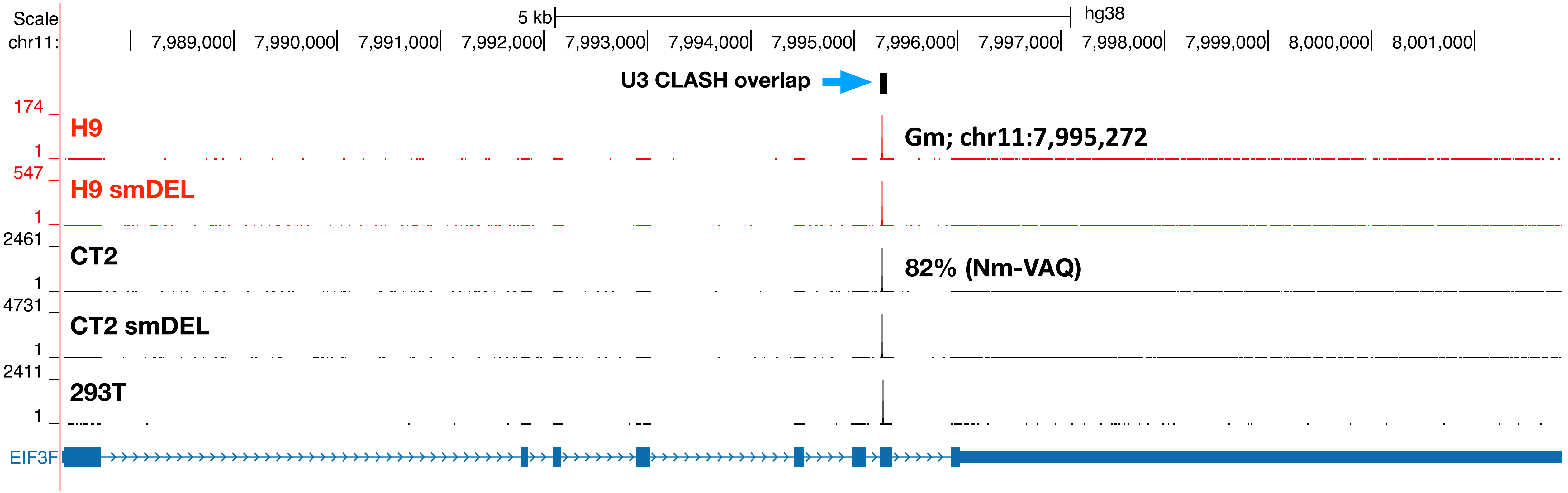

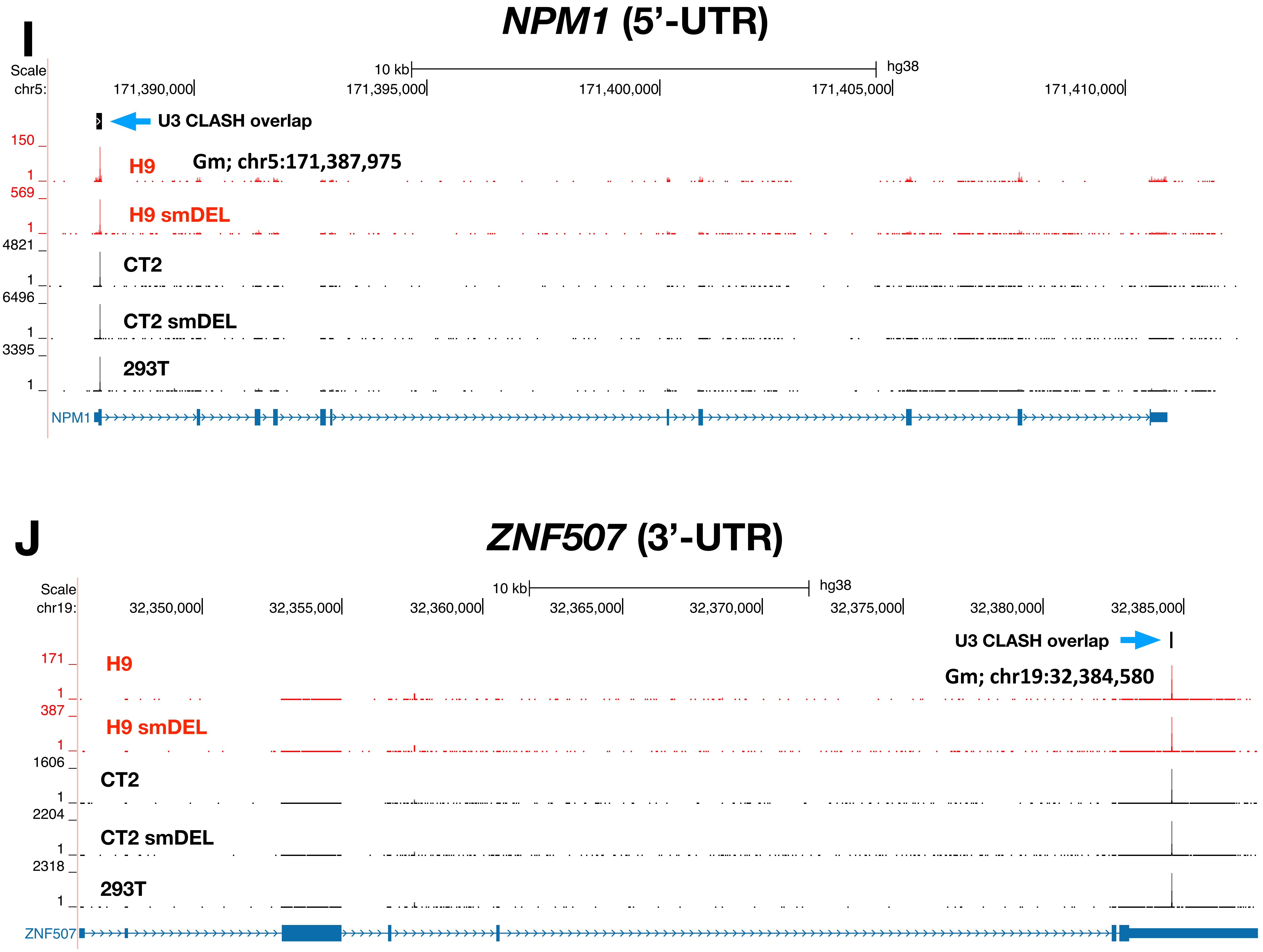

K

#### *FAM177B* (3'-UTR)

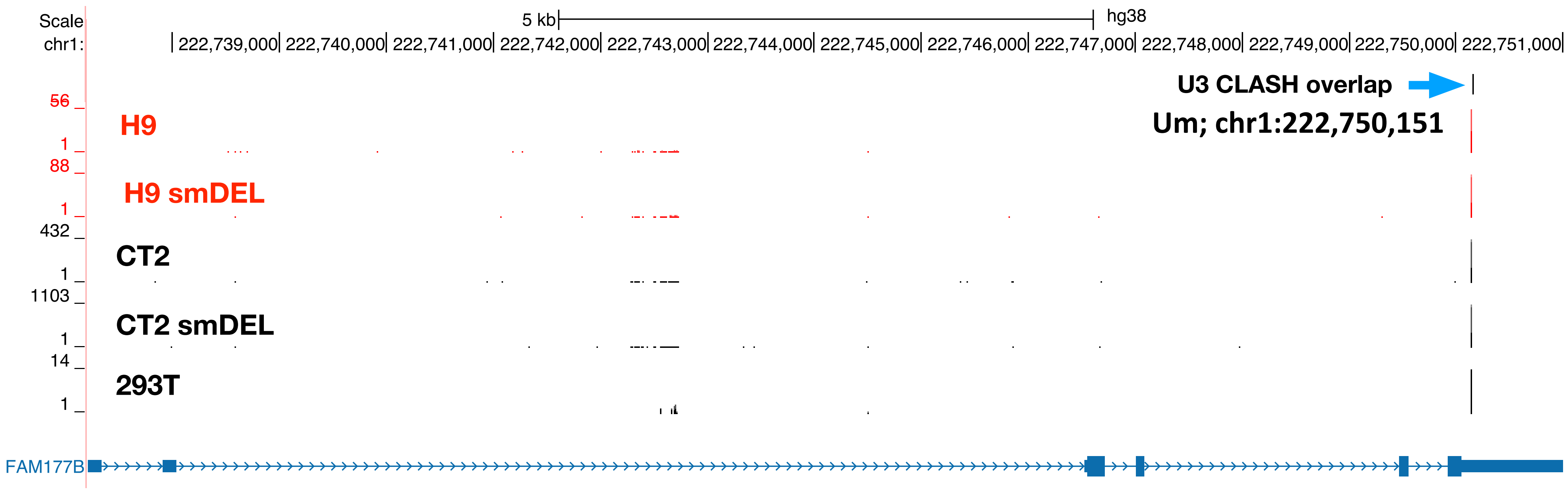


L

#### *SYNJ2BP* (3'-UTR)

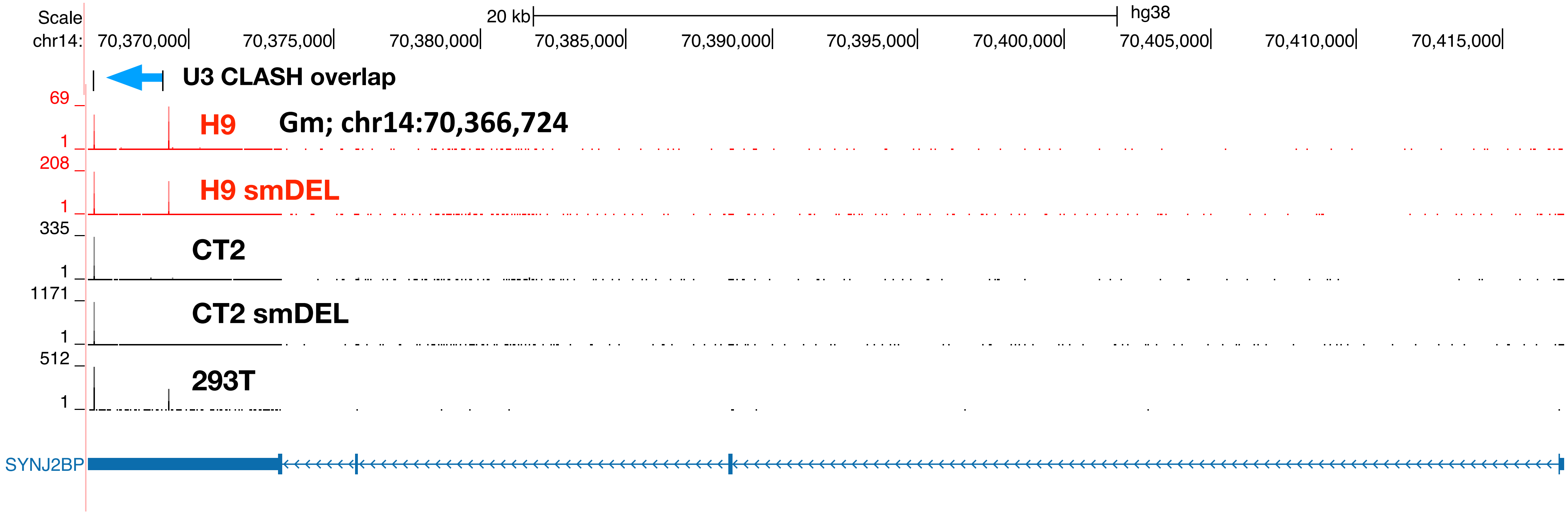

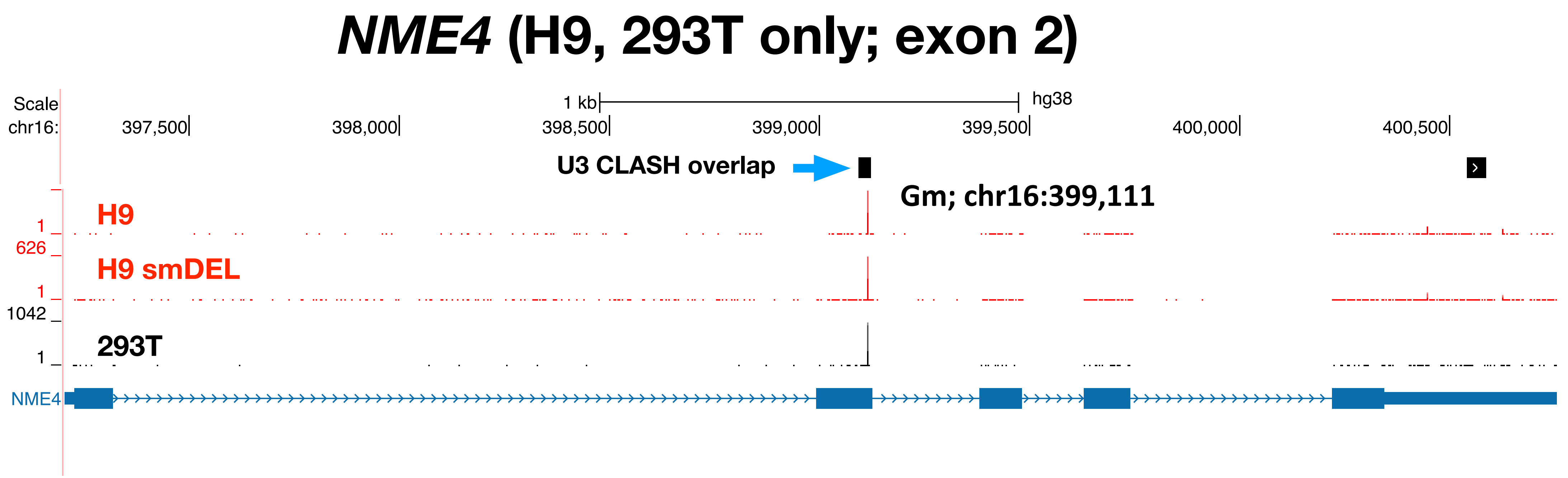

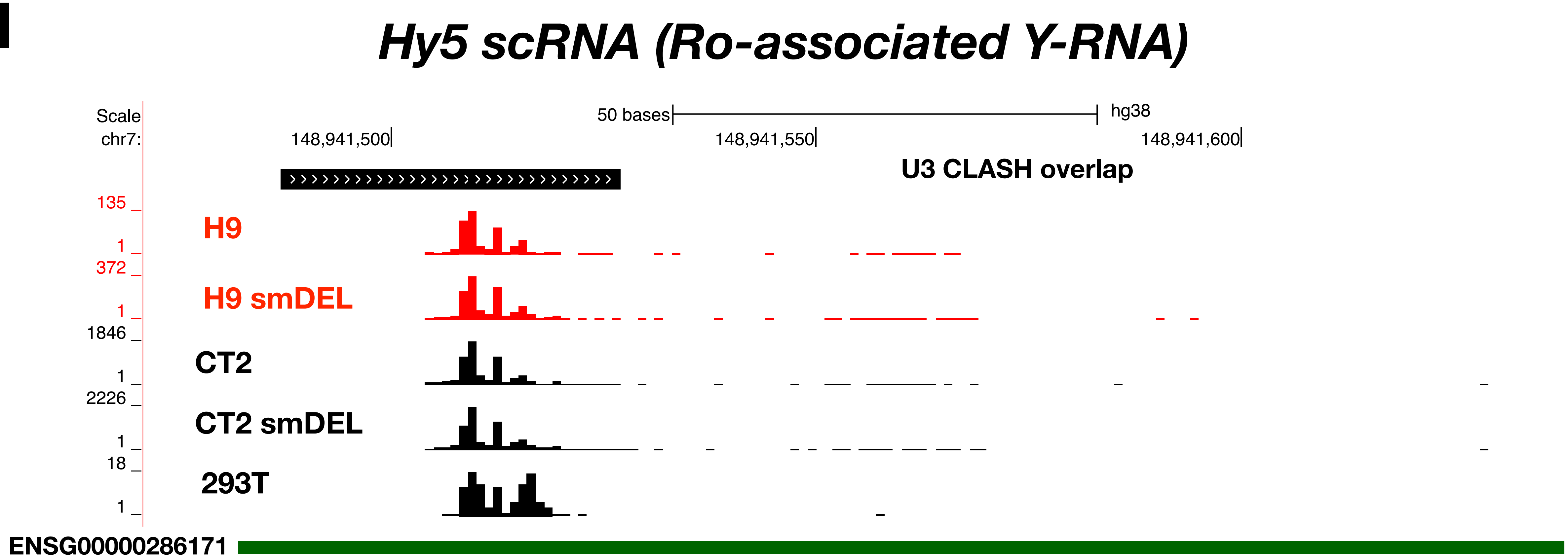
