## Supplemental File 4 for "Genome-wide profiling of RNA 2’-*O*-methylation in neurons and identification of orphan snoRNA targets"

**Nm-VAQ DNA-RNA Chimeras**

| **Chimera** | **Sequences 5’->3’** |
| --- | --- |
| 18S-G1328 | CCACmCmAmCmCmCmAmCmGmGmAmAmUmCmG |
| 18S-A159 | TTACmCmAmCmAmGmUmUmAmUmCmCmAmAmG |
| 18S-U1445 | AAGAmAmGmUmUmGmGmGmGmGmAmCmGmC |
| 28S-Ψ3818 | ATTCmAmUmGmCmGmCmGmUmCmAmCmUmA |
| ACTG1 | TGCGmGmAmUmGmUmCmCmAmCmGmUmCmA |
| ACTB-2 | GCGGmAmAmCmCmGmCmUmCmAmUmUmGmC |
| SNCG | CGCTmGmCmUmCmAmCmCmAmCmAmGmCmC |
| GUK1 | CCGAmGmAmAmCmUmCmGmGmCmAmUmGmC |
| NEFM | CGCGmCmAmAmCmCmGmCmGmCmCmUmCmC |
| MIAT | GAACmCmAmCmAmUmGmGmGmUmGmGmAmC |
| TPT1 | GACGmAmCmGmGmCmGmCmUmAmGmCmUmU |
| EIF3F | CAGCmUmGmAmCmAmCmCmUmUmUmCmCmA |
| RN7SK | CTTGmAmGmAmGmCmUmUmGmUmUmUmGmG |
| AZIN1 | CAGGmGmUmAmGmUmGmCmCmAmAmAmCmU |
| TKT | CGGGmUmCmAmCmUmCmUmGmGmCmCmCmA |
| NUDT21 | GGGCmUmUmCmGmUmCmUmGmCmUmGmGmA |
| FAM171A1 | CTCCmAmUmCmCmCmUmGmUmGmGmCmAmC |
| FGF13 | GACTmUmCmCmUmUmAmAmCmAmAmAmGmC |
| C12orf57 | GAACmAmUmCmAmAmGmGmCmCmUmUmUmC |
| MT-CO1 | AGGAmUmGmUmUmUmCmAmUmGmUmGmGmU |

**qPCR Primers**

| **Target** | **Forward 5’->3’** | **Reverse 5’->3’** |
| --- | --- | --- |
| 18S-G1328 | CTCAACACGGGAAACCTCAC | TCGGAATTAACCAGACAAATCGC |
| 18S-A159/U161 | CCGGTACAGTGAAACTGCGA | CTGATAAATGCACGCATCCCC |
| 18S-U1445 | TAGTTACGCGACCCCCG | GGGCATCACAGACCTGT |
| 28S-Ψ3818 | GCCAAATGCCTCGTCATC | CCGCTGATTCCGCCAAG |
| ACTG1 | GTATGGAATCTTGCGGCATC | CCTTGATCTTCATGGTGCTG |
| ACTB-1 | GCATGGAGTCCTGTGGCATC | TTGATCTTCATTGTGCTGGG |
| ACTB-2 | CCTGGAGAAGAGCTACGAGC | CCAGGAAGGAAGGCTGGAAG |
| SNCG | GTGGCCGAGAAGACCAAGG | CTTGCGCACCACCCCGGAG |
| GUK1 | TTCATCGAGCATGCCGAGTT | CAGATCGGTGGCCTTGATGT |
| NEFM | AAGACATCCACCGGCTCAAG | TTGTCCAGCTCCACCTTGAC |
| MIAT | GAGGCATCTGTCCACCCATG | AGAGGGAGGATCACCTTGGT |
| TPT1 | TCCCCCCGAGCGCCGCTC | GGCTGATGAGGTCCCGGTA |
| EIF3F | AGCTGACAATACTGTGGGCC | GAGGTGGGGGAAGAAACCAG |
| RN7SK | TAGAGGAGGACCGGTCTTCG | GCGCCTCATTTGGATGTGTC |
| AZIN1 | CTCTGCCCCAATCATCTCCC | AGCACATTCCAAGAGATGCCT |
| TKT | CACCGCTTCTGGTAGATG | ACCTTGTGGCCATTCTAG |
| NUTD21 | GGGGTCACTCAGTTCGG | AGGTTGATGGTGCGCTCC |
| FAM171A1 | TAGCGAAGATAGACTAGAG | GCTCGCTTGCTCAAACC |
| FGF13 | TCCTCTCTCTCTGTGTCTGCT | TGTCCTCATCTTTGGTGCCA |
| C12orf57 | CTGAAGGCGCTGTTTCTGC | CTAAGTCACAGGCGCATTTA |
| MT-CO1 | AACACTTTCTCGGCCTATCC | TTTTCGCTTCGAAGCGAAGG |
